# An amino residue that guides the correct photoassembly of the water-oxidation complex but is not required for high affinity Mn^2+^ binding

**DOI:** 10.1101/2021.11.29.470031

**Authors:** Anton P. Avramov, Minquan Zhang, Robert L. Burnap

## Abstract

The assembly of the Mn_4_O_5_Ca cluster of the photosystem II (PSII) starts from the initial binding and photooxidation of the first Mn^2+^ at a high affinity site (HAS). Recent cryo-EM apo-PSII structures reveal an altered geometry of amino ligands in this region and suggest the involvement of D1-Glu189 ligand in the formation of the HAS. We now find that Gln and Lys substitution mutants photoactivate with reduced quantum efficiency compared to the wild-type. However, the affinity of Mn^2+^ at the HAS in D1-E189K was very similar to the wild-type (~2.2 μM). Thus, we conclude that D1-E189 does not form the HAS (~2.9 μM) and that the reduced quantum efficiency of photoactivation in D1-E189K cannot be ascribed to the initial photooxidation of Mn^2+^ at the HAS. Besides reduced quantum efficiency, the D1-E189K mutant exhibits a large fraction of centers that fail to recover activity during photoactivation starting early in the assembly phase, becoming recalcitrant to further assembly. Fluorescence relaxation kinetics indicate on the presence of an alternative route for the charge recombination in Mn-depleted samples in all studied mutants and exclude damage to the photochemical reaction center as the cause for the recalcitrant centers failing to assemble and show that dark incubation of cells reverses some of the inactivation. This reversibility would explain the ability of these mutants to accumulate a significant fraction of active PSII during extended periods of cell growth. The failed recovery in the fraction of inactive centers appears to a reversible mis-assembly involving the accumulation of photooxidized, but non-catalytic high valence Mn at the donor side of photosystem II, and that a reductive mechanism exists for restoration of assembly capacity at sites incurring mis-assembly. Given the established role of Ca^2+^ in preventing misassembled Mn, we conclude that D1-E189K mutant impairs the ligation of Ca^2+^ at its effector site in all PSII centers that consequently leads to the mis-assembly resulting in accumulation of non-catalytic Mn at the donor side of PSII. Our data indicate that D1-E189 is not functionally involved in Mn^2+^ oxidation\binding at the HAS but rather involved in Ca^2+^ ligation and steps following the initial Mn^2+^ photooxidation.

## 1. Introduction

The photosystem II (PSII) reaction center complex of photosynthetic organisms catalyzes the extraction of electrons from water molecules to plastoquinone, harnessing light as the energy source to drive this highly endergonic electron transfer. Primary charge separation in PSII involves the formation of the highly oxidized electron donor P_680_^+^ and a pheophytin within the D1/D2 polypeptide core buried within the large PSII complex. P_680_^+^ is a powerful oxidant capable of extracting the electrons from the H_2_O-oxidizing Mn_4_O_5_Ca metal cluster via a neighboring redox active tyrosine residue (Y_Z_) of the D1 protein (1, 2). After each charge separation, the Mn cluster advances from a more reduced to a more oxidized state, with every state corresponding to one of the five so-called storage states (S_0_, S_1_, S_2_, S_3_, and S_4_) of S-cycle. The oxidation of two H_2_O molecules H_2_O with the release of molecular oxygen is triggered by the formation of S_4_ state, which is rapidly reduced by the substrate H_2_O to the S_0_ state, representing Mn cluster ground state (3).

The assembly of the Mn_4_O_5_Ca metal cluster within the protein matrix of the water oxidizing complex (WOC) is termed as photoactivation. It is an oxidative process that stepwise incorporates Mn^2+^ into the high valent Mn^≥3+^ oxo-bridged structure that characterizes the H_2_O-oxidation catalyst. Importantly, the first two Mn^2+^ oxidation events are separated by a poorly understood rate-limiting light independent process termed as dark rearrangement, k_A_ (Scheme 1).

**Scheme 1.**
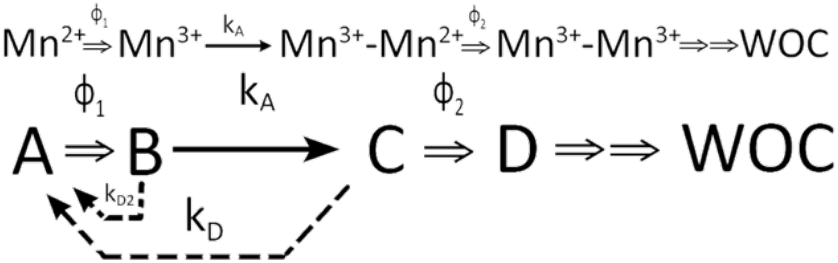
Kinetic scheme of basic two-quantum mechanism of PSII photoactivation. Double arrows indicate light-activated processes with the quantum efficiencies ***ϕ***_1_ and ***ϕ***_2_ representing the first and second photooxidative events in the assembly sequence, *k_A_* representing the “dark” rearrangement, and *k*_D_, representing the decay of intermediates. After the initial two Mn are photoligated, subsequent Mn appear to be added with high quantum yield.

Remarkably, photoactivation utilizes essentially the same redox active cofactors as the regular electron transfer pathway during H_2_O-oxidation catalysis. It begins with photochemical charge separation with the D1/D2 reaction center core with the formation of the highly oxidizing P_680_^+^ species. During assembly, P_680_^+^ extracts electrons, again via Y_z_, from Mn^2+^ ions that diffuse from the soluble phase and bind within the apo-WOC at the so-called high affinity site (HAS). These Mn^2+^ ions successively become photooxidatively incorporated into the growing metal cluster [reviewed in (4)]. Current evidence suggests that Mn^2+^ photooxidation corresponds to the formation of a Mn^2+^ hydroxide intermediate at the HAS (5, 6). The exact location of the HAS is still a matter of debate. Surveys of amino acids at the PSII donor side using site-directed mutagenesis revealed D1-Asp170 is most important for the high affinity Mn^2+^ binding and photooxidation in apo-PSII (7–11). Not surprisingly, the affinity for Mn^2+^ ions (K_M_ < 10 μM) at the HAS (5, 8, 12) has strong pH dependence, highlighting with the importance of the protonation state of the amino acid residues coordinating Mn^2+^ and the likely hydroxide form of the bound Mn^2+^ prior to photooxidation (5). In the crystal structure of the fully assembled complex, D1-Asp170 provides one of two oppositely arranged axial monodentate carboxyl ligands of the so-called dangler Mn (Mn4) in intact WOC. Interestingly, the other ligand, D1-Glu333, does not seem to be directly involved in the formation of the HAS of apo-PSII, although it weakly influences the affinity of Mn^2+^ at the HAS (11, 13, 14). The evidence that D1-Glu333 does not partake in the formation of the HAS is consistent with recent cryo-EM structures of apo-PSII revealing that the carboxyl domain of the D1 protein adopts a very different conformation and with several residues, including D1-Glu333, are shifted several Ångstroms away from their final positions in the assembled complex (15, 16). The absence of direct Mn coordination by D1-Glu333 is interesting in the light of the assumption that the high affinity binding of Mn^2+^ ion requires two carboxylate moieties (17). Interestingly, D1-His332 is slightly shifted towards hypothetical HAS in apo-PSII and could be involved in coordination of the Mn^2+^ ion (15, 16). The closest carboxylate moiety in apo-PSII to D1-Asp170 is D1-Glu189, which occupies a similar position (1.1 Å difference relative to the assembled structure) in the presence and absence of the Mn cluster. The distance between D1-Glu189 and D1-Asp170 in apo-PSII is ~6.0 Å, which is in agreement with the distances found in mature PSII between carboxylate and intervening Mn ions and, on this basis, has been proposed that D1-Glu189 and D1-Asp170 participate in the formation of the HAS, perhaps in conjunction with D1-His332 (15, 16). Importantly, the most recent cryo-EM structures (16) of apo-PSII reveal an unidentified a heavy atom, probably either Mn or Ca, located midway between the carboxylate moieties of D1-Glu189 and D1-Asp170. This precise spatial arrangement is not found in the assembled complex since these residues are slightly shifted in orientation compared to their assembled configuration and the heavy atom appears to be in slightly different place than any of the metal ions of the assembled Mn_4_O_5_Ca. The D1-Glu189 and D1-Asp170 residues share another important characteristic, at least in the assembled complex: besides providing monodentate coordination to Mn (Mn1 and Mn4, respectively), their carboxylate groups each provide a second monodentate ligand to the sole Ca ion of the Mn_4_O_5_Ca (14). In this context, the photoactivation of the Mn_4_O_5_Ca requires the presence of Ca^2+^ during the assembly process and without Ca^2+^, inactive high valent Mn accumulates (18). Previous work has shown that binding of Ca^2+^ to the Ca^2+^ effector site (CES) during photoactivation is essential to avoid the competitive binding of Mn^2+^ to the CES resulting in this inactivation (19). Additionally, binding of Ca^2+^ to the CES stabilizes the intermediates of photoactivation, although excessive [Ca^2+^] diminishes the quantum efficiency of photoactivation by binding to a second Mn^2+^ binding site.

Besides providing a ligand to Mn1, D1-Glu189 modulates the redox properties of Y_Z_ in addition to those of the Mn cluster (20, 21). Additionally, D1-Glu189 is involved in the formation of a H-bonding network, potentially accepting a proton from D1-His190 (22, 23). Most of the D1-Glu189 site-directed mutants assemble partial or complete Mn clusters, possibly having defects in cluster assembly (20, 24). Most of these mutants have lost the ability to evolve oxygen, despite containing photooxidizable Mn *in vivo* (20). Recent time-resolved X-ray crystallography indicates that D1-Glu189 side chain moves away a from the Mn_4_O_5_Ca during the S_2_-S_3_ transition, such that the ligand to the Ca ion is broken, but the D1-Glu189-Mn1 ligation is maintained (25, 26). This movement appears to facilitate the Ca^2+^-mediated insertion of substrate H_2_O between Mn1 and the Ca, thereby producing the O_x_ (O_6_) proposed to be one of the two oxygen atoms of the O_2_ formed in the final stages of water oxidation (25–28). On the other hand, the fact that several mutations with disparate chemical properties can substitute for glutamate suggests this role is not decisive in the catalytic mechanism.

Given recent results indicating that the D1-Glu189 is potentially involved in the formation of the HAS of PSII, we studied the photoactivation characteristics of three site directed mutants with lysine, arginine and glutamine substitutions at the 189 position of D1 protein, which are substitutions that permit relatively high rates of oxygen evolution (20), yet are shown to alter the assembly if the Mn_4_O_5_Ca.

## 2. Materials and Methods

### 2.1. Strains and Growth Conditions

The glucose-tolerant strain of *Synechocystis sp*. PCC6803 GT (hereafter *Synechocystis*) (29) served as the basis for all strains conducted in this study. The wild-type control strain (WT control) was constructed by transforming strain 4E3 HT-3 returning the wild-type *psbA2* gene to its native location and with its native promoter using *psbA2* using in the suicide plasmid pRD1031 as described previously (30). The 4E3 HT-3 strain lacks all three *psbA* genes and has the CP47 gene hexahistadine tag at its carboxy terminus. Transformation of 4E3 HT-3 with the wild-type *psbA2* gene restored wild-type levels of D1 expression and oxygen evolution under normal growth conditions (30). Site-directed mutations in the *psbA2* gene encoding the D1-E189Q, D1-E189K and D1-E189R amino acid substitutions involved similar transformations, but using site-directed mutant *psbA2* alleles resulting the mutant strains (31). All *Synechocystis* strains were routinely maintained on BG-11 agar plates with addition of 10 μM DCMU and 5mM glucose as described previously, to prevent reversion and suppressor mutations since these conditions remove any selective advantage higher PSII activity (32). Experimental cultures were grown in 1 L flat flasks in 800 mL of BG-11 with the addition of 10 mM HEPES-NaOH pH 8.0 and 5mM glucose under PFD (photon flux density) of ~100 μmol m^−2^ s^−1^ at 30 °C. Light intensity was measured with Walz light meter (Germany). All experimental cultures were harvested in late log phase (O.D. 750nm ~1.2-1.5) and checking the variable fluorescence (F_V_) value maximal for every strain (supplementary Table 1) as determined with a PSI fluorometer (PSI instruments) using F_v_ = (F_max_ – F_0_)/ F_0_.

### 2.2. Hydroxylamine Extraction of Cells

Cultures were centrifuged in 1 L bottles at 6000g (Sorvall, F9 rotor) for 12 minutes at room temperature. Extraction of the Mn_4_O_5_Ca using hydroxylamine (HA) was performed as described previously (33, 34). Briefly, cells were washed in BG-11 and concentrated to a chlorophyll concentration of 100 μg/mL, HA was added to a cell suspension from a 400 mM freshly prepared stock to a final concentration of 10mM (35). Cell suspension was incubated in the dark on a rotary shaker at ~120 rpm at room temperature for 12 minutes. After the incubation, cells were diluted with 5x volume of BG-11 and centrifuged at 10200g (Sorvall, F14 rotor) at room temperature for 5 minutes. This washing step was repeated for 7 more times. After the washing was complete, cells were resuspended to [Chl] of 100 μg/mL and kept in the dark with shaking at ~120 rpm.

### 2.3. Photoactivation of HA-Extracted cells

HA-extracted were either photoactivated using single-turnover Xenon lamp flashes or continuous light (Cool White fluorescent) at an intensity of ~50 μmol m^−2^ s^−1^. Single-turnover Xenon lamp flash illumination was carried out as described previously in (34). A 400 μL aliquot of cells containing ~40 μg of chlorophyll was taken from the cell suspension kept in the dark. Cells were placed on a sample vessel fashioned out of an aluminum weighing cup with the stirring bar and covered with yellow UV filter to minimize UV damage to the sample. The sample was subjected to saturating single-turnover Xe flashes to promote photoactivation. Saturation of samples by Xe flashes was ensured by pre-tests using neutral density filters. Flash illumination was followed by the measurement of light-saturated rates of O_2_ evolution to evaluate restoration of catalytic activity. To accomplish this, aliquots containing 10 μg of Chl were withdrawn from the photoactivated cell suspension and placed in the Clark-type electrode (Yellow Springs Instruments) chamber containing buffered BG-11 with the addition of 1mM DCBQ and 1mM potassium ferricyanide. The oxygen evolution rate was measured in response to saturating orange light (>570 nm) at 30°C.

### 2.4. Curve fitting and data analysis

Experiments measuring the photoactivation of the Mn-depleted PSII as a function of single-turnover flashes and a function of the flash interval allowed us to evaluate both the quantum efficiency of photoactivation and to estimate the kinetic parameters for the dark rearrangement and decay of intermediates (36). The rate constants for the dark rearrangement, *k*_A_, and the decay of intermediates, *k*_D_, were determined by deriving kinetic parameters from the rising and falling slopes of the bell-shaped curve in plots of photoactivation as a function of the flash interval curve fitted to the equation (35, 37–39):

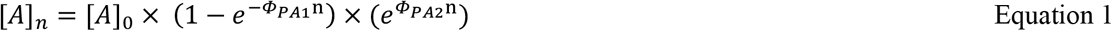

Equation 1 Where [A]_n_ represents the yield of active centers on the *n*th flash, [A]_0_ is the concentration of apo-PSII centers prior to the photoactivation, ***Φ**_PA1_*, and ***Φ**_PA2_* represent two apparent phases of photoactivation: a more efficient phase most obvious in the beginning of the flash sequence and lower efficiency phase afterwards.

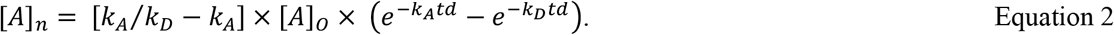

Where *k*_A_ and *k*_D_ represents for the dark rearrangement and the decay of intermediates, respectively.

### 2.5. Fluorescence relaxation kinetics

Flash-induced fluorescence yield relaxation kinetics was measured as described by Vass and coworkers (40) using FL-3300 fluorometer (PSI Instruments, Czech Republic). The duration of actinic and measuring light pulses were 30 μs and 9 μs, respectively, and relaxation of fluorescence yield was measured from range of 100 μs to 100 s. Samples were diluted to a concentration of 5 μg of Chl/mL in 3 mL of BG-11 and placed into the 4-sided clear cuvette for measurement. Multi component fluorescence relaxation analysis was performed as earlier by Vass (40, 41) by using fitting function with two or three components:

## 3. Results

### 3.1. Photoactivation in continuous light shows altered Mn_4_O_5_Ca assembly in mutants

There is consensus that D1-Asp170 is critical for the binding and photooxidation of Mn^2+^ at the HAS (7, 8, 11, 35), consistent with it providing a monodentate ligand to Mn4 in the assembled complex (14). However, recent cryo-EM studies of Mn-depleted PSII and PSII assembly intermediates that lack the Mn_4_O_5_Ca indicate the HAS is structurally quite different in the apo-PSII and identify two additional residues potentially involved in the formation of the HAS in apo-PSII and thus potentially involved in photoactivation. These residues, D1-Glu189 and D1-His332, are shifted towards the D1-Asp170 side chain compared to mature PSII. Their close proximity to D1-Asp170 potentially allows for Mn^2+^ to be coordinated between them, which is in accord with the observation of an unidentified heavy atom located midway between the D1-Asp170 and D1-Glu189 carboxyl groups (15, 16). To test the general hypothesis that D1-Glu189 modulates photoactivation, two amino acid substitution mutants capable of relatively high rates of oxygen evolution, D1-E189Q, D1-E189K and D1-E189R (**Supplementary material**) were investigated. In addition, the D1-D170E mutant, which was previously shown to have a lower quantum efficiency of photoactivation (35) was also included in this study for comparison.

The oxygen evolution rates in freshly isolated cells are shown in Figure 1A. The site-directed mutants exhibit rates that are approximately 60% of the WT control, with the exception of D1-E189K which was about 80% (20). The results confirm the initial studies on these mutants and demonstrate that even relatively drastic amino acid substitutions such as D1-E189K still allow the assembly of active PSII under the protracted conditions of growth and that the assembled Mn_4_O_5_Ca has robust H_2_O-oxidation activity despite the mutations. To gain information on whether the mutations affect the kinetics of assembly, cells were treated with hydroxylamine to remove active Mn and samples were photoactivated under continuous light (**Fig. 1B**). The extent of recovered O_2_ evolution activity is expressed as a ratio of the recovered rate to the rate prior to HA-extraction. Consistent with previous results, WT control cells recovered >90% of their original activity within 15 min of illumination. Photoactivation of HA-extracted D1-D170E cells restored approximately 70% of the original activity, but this took longer, ~40 minutes. This consistent with the flash experiments (next section) and previous findings (35) showing reduced quantum efficiency of photoactivation. Thus, substitution of Asp170 of the HAS, with the slightly large Glu side chain, permits assembly, but with reduced quantum efficiency. Consistent with the nature of the amino acid substitutions, the restoration of the O_2_ activity was dramatically different D1-E189K and D1-E189Q mutants: D1-E189Q was able to show similar restoration level to D1-D170E. However, it required only 20 minutes to reach 70% of its initial activity, while D1-E189K with the highest net O_2_ evolution activity restored only 40% even after 40 minutes of illumination respectively. Such low level of restoration in D1-E189K despites the relatively high extent of assembly possible when grown in culture.

**Figure 1.**
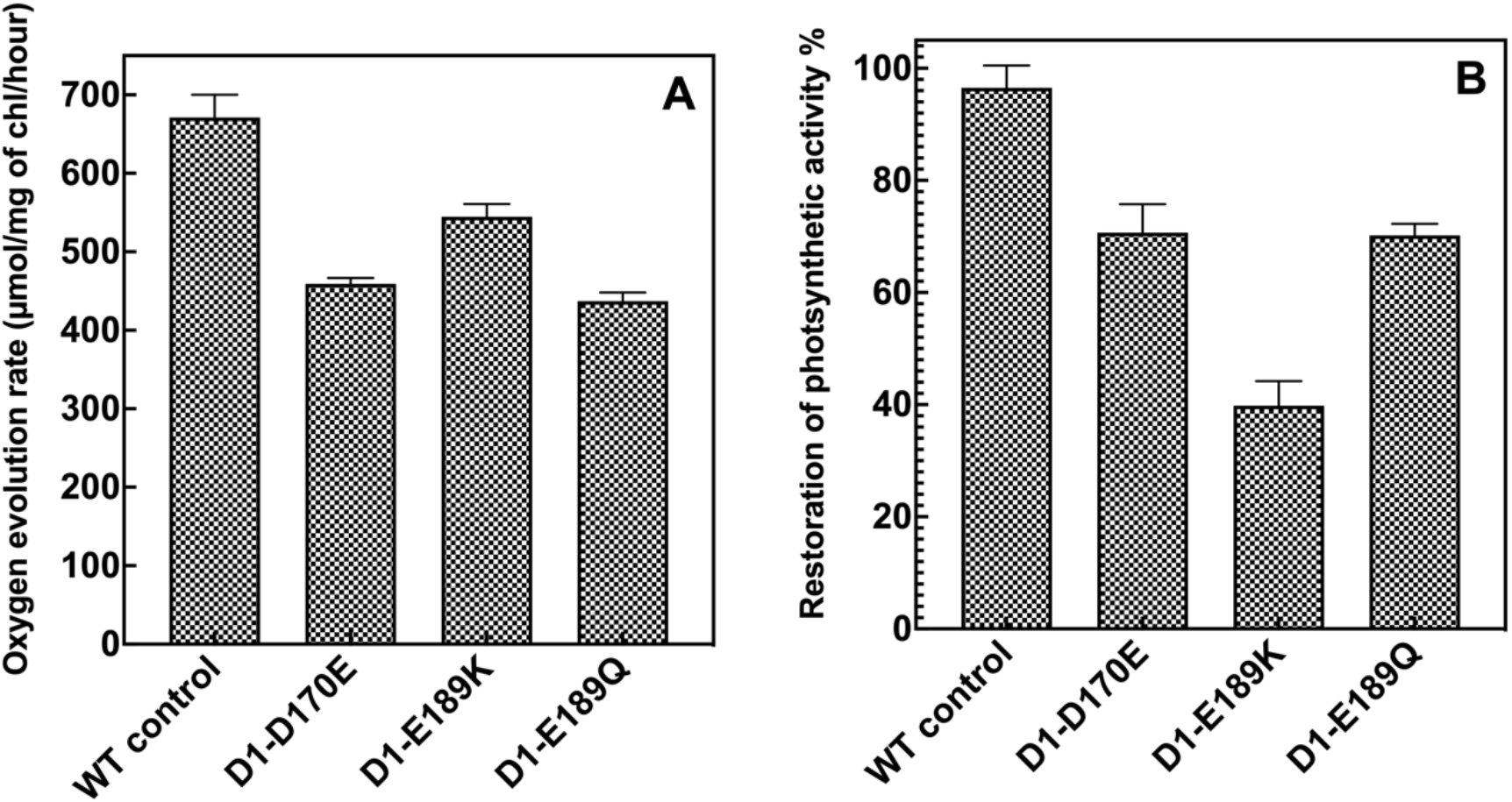
Light-saturated O_2_-evolution activity of *Synechocystis* cells (A) harvested directly from growth culture. (B) Recovery of light-saturated O_2_-evolution activity of HA-treated *Synechocystis* cells after photoactivation under continuous illumination. HA-treated cells were exposed to continuous illumination (~20 μmol m^−2^ s^−1^ PD) while being gently shaken. Oxygen evolution activity was measured using a Clark-type electrode in HN buffer (10mM HEPES, 30mM NaCl, pH 7.2 in the presence of 1mM DCBQ and 1mM potassium ferricyanide). Oxygen evolution was measures in response to saturating (>570 nm) illumination at 30°C. The original light-saturated O_2_-evolution rates prior HA extraction were 671 ± 29, 459 ± 7, 544 ± 16, and 437 ± 11 μmol of O_2_ (mg Chl)^−1^ h^−1^, for WT control, D1-D170E, D1-E189K and D1-E189Q respectively. Error bars represent SD with n ≥ 3.

### 3.2. Restoration of the O_2_ evolving activity

The photoactivation as a function of the number (**Fig. 2**) and frequency (**Supplementary results**) of saturating, single-turnover Xe flashes allows us to gather the values of quantum efficiency of photoactivation i.e., estimate values that reflect the per flash development of O_2_ evolution activity in Mn-depleted samples and estimate the stability of the intermediates formed during the photoactivation.

**Figure 2.**
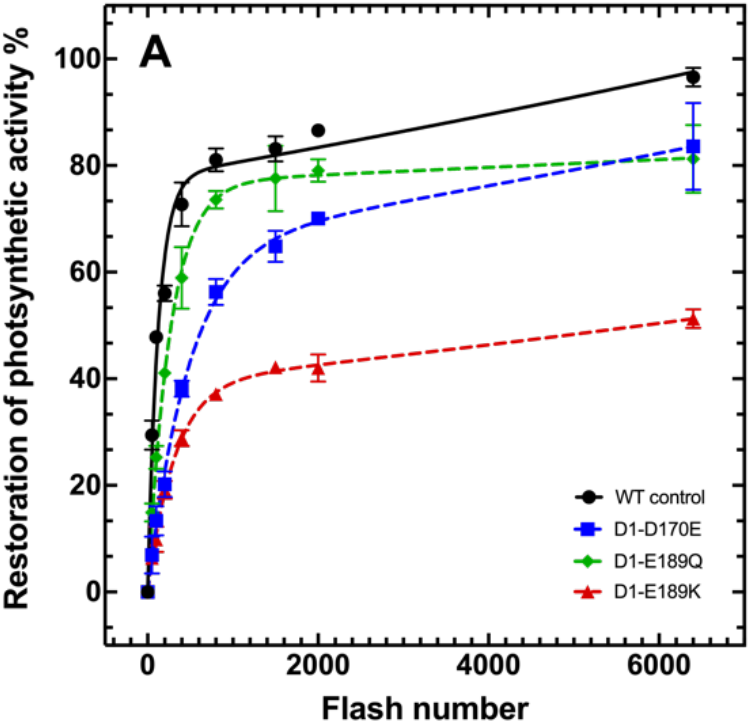
Photoactivation of HA-extracted cells as a function of the single-turnover flash number. Series of xenon lamp flashes were given at uniform frequency of 2 Hz (500ms flash interval) to hydroxylamine-treated WT control, D1-D170E, D1-E189Q, D1-E189K,. The percentage of O_2_ evolution recovery was calculated as a fraction from the O_2_ evolution rate prior HA extraction as described in Figure 1. Error bars represent SD with n ≥ 3.

The yields of photoactivation as a function of flash number are illustrated in Figure 2. For this experiment, the uniform flash interval of 500ms (2 Hz) was used to photoactivate the samples. Consistent with the previous results (34, 35, 42), HA-treated WT control cells reached approximately 85-90% if initial activity after 2000 flashes, while 6400 flashes resulted in almost complete restoration of O_2_ activity (**Fig 2**). The D1-E189Q mutant exhibits similarly shaped photoactivation curve as the WT control, however it never approaches complete photoactivation. The maximum restoration of O_2_ activity of 80% was observed after 2000 single-turnover flashes and percent activity does not significantly increase even after 6400 flashes. This behavior contrasts with both WT-control, which is capable of reaching nearly 100% of initial activity, and D1-D170E mutant that exhibits lower quantum efficiency of photoactivation but nevertheless continues to develop O_2_ evolution rate throughout the entire series of photoactivation flashes although not reaching levels as high as the WT even after 6400 flashes. Unlike D1-E189Q, D1-E189K caused a severe inhibition of photoactivation under flash illumination (**Figs. 1B, 2**) even though the maximal O_2_ net rates prior to extraction being similar to the other mutants (**Fig 1A**). A series of 2000 photoactivation flashes given to D1-E189K mutants resulted only in 40% interestingly, the level of photoactivation shows little increase even after 6400 flashes.

To derive the constants of both high and low efficiency phases of photoactivation seen in **Fig 2**, the obtained data was fit into the double exponent equation 1. The quantum efficiency of the photoactivation, ***Φ**_PA1,2_* (**Table 2**), in all studied mutants indicates the severe effect of the mutations at the HAS on the overall efficiency of the light driven assembly of the Mn_4_O_5_Ca cluster but distributed differently between the two phases. Interestingly D1-D170E mutant, which is shown to have a significantly lower quantum efficiency of photoactivation is capable of high levels of restoration. Although D1-E189K show higher ***Φ**_PA1_* than D1-D170E, it fails to reassemble Mn cluster in the majority of the PSII centers indicating on an irreversible inhibition of the photoactivation. All of the studied D1-E189 mutants share a similar feature of an efficient initial photoactivation phase within first 400-800 flashes followed by a significant decrease in the efficiency in assembly and failure to restore activity in a large fraction of centers.

**Table 1.** Multicomponent kinetic equations used to fit the data from the data plots obtained from fluorescence relaxation kinetics experiments.

| Components | Equation |
| --- | --- |
| Fast – Exponential<br>Middle – Exponential<br>Slow – Hyperbolic | $F(t) - F_0 = A_1 \exp\left(-\frac{t}{T_1}\right) + A_2 \exp\left(-\frac{t}{T_2}\right) + A_3/(1 + \frac{t}{T_3})$ |
| Fast – Exponential<br>Slow – Hyperbolic | $F(t) - F_0 = A_1 \exp\left(-\frac{t}{T_1}\right) + A_2/(1 + \frac{t}{T_2})$ |
| Fast – Exponential<br>Middle – Hyperbolic<br>Slow – Hyperbolic | $F(t) - F_0 = A_1 \exp\left(-\frac{t}{T_1}\right) + A_2/(1 + \frac{t}{T_2}) + A_3/(1 + \frac{t}{T_3})$ |

Where $F(t) - F_0$ is relative fluorescence yield, $A_1$ - $A_3$ are decay amplitudes, $T_1$ - $T_3$ are time constants from which the half-times can be calculated via $t_{1/2} = \ln 2 T$ for exponential components and $t_{1/2} = T$ for hyperbolic component.

**Table 2.** Quantum efficiency of photoactivation of WT control, D1-D170E, D1-E189K and D1-E189Q mutant strains. Kinetic fit parameters of ***Φ**_PA1_* and ***Φ**_PA2_* obtained from the data from the plots in Figure 2 was fit into Equation 1.

| Strain | Quantum efficiency of Photoactivation, $\Phi_{PA1}$ | Quantum efficiency Restoration, $\Phi_{PA2}$ | R |
| --- | --- | --- | --- |
| WT control | $8.1 \times 10^{-3}$ | $3.6 \times 10^{-5}$ | 0.98 |
| D1-D170E | $2.1 \times 10^{-3}$ | $3.8 \times 10^{-5}$ | 0.99 |
| D1-E189K | $3.2 \times 10^{-3}$ | $4.2 \times 10^{-5}$ | 0.99 |
| D1-E189Q | $3.8 \times 10^{-3}$ | $9.1 \times 10^{-6}$ | 0.99 |

### 3.3. Mutations at D1-E189 does not alter the affinity at the HAS

Among the studied D1-E189 site directed mutants, D1-E189K represents greater interest due to the intriguing relationship between high net oxygen evolution activity (**Fig. 1A**), an impaired ability for *de novo* assembly of the Mn_4_CaO_5_ cluster (**Figs. 1B and 2**), and no apparent changes to the dark rearrangement and stability of photoactivation intermediates (**Fig. S1, Table S2**). While the photoactivation experiments fail to explain intriguing behavior of D1-E189 mutants, fluorescence relaxation kinetics experiments (**Supplementary results**) potentially indicate on a lower Mn^2+^ binding affinity at the HAS (**Fig S2, Table S2**). To estimate the effect of the lysin substitution on the Mn^2+^ binding at the HAS during photoactivation, the electron donation from Mn^2+^ to Y_z_ was measured in highly purified Mn-depleted PSII core complexes as a function variable fluorescence (43). Mn^2+^ binding constant at the HAS in Mn-depleted PSII was earlier estimated at approximately 1μM and has been shown to be strong pH dependent (5, 8, 44, 45) (46). When the first charge separation occur, the redox-active Y_z_ re-reduces a primary chlorophyll donor, P_680_^+^. At the time of the second charge separation in the Mn-depleted PSII, Y_z_ is already in the oxidized state and in the absence of electron donor it is unable to re-reduce P_680_^+^, which results in the strong variable fluorescence quenching by meta-stable P_680_^+^. In the absence of Mn^2+^ bound at the HAS, second and subsequent charge separations result in essentially complete fluorescence quenching, however binding of Mn^2+^ at the HAS can completely restores electron transport, yielding high fluorescence after each flash (5, 43, 47).

The electron flow from Mn^2+^ to Y_Z_ has a strong [Mn^2+^] dependence (**Fig. S4, Table 4**), at very low Mn^2+^ concentrations (<1μM) Y_Z_ is not able to re-reduce the entire population of P_680_^+^ resulting in fluorescence quenching, however at K_m_ and saturating [Mn^2+^] the reduction of P_680_^+^ produce a high fluorescence yield allowing to estimate the binding affinity of Mn^2+^ at the HAS as a function of variable fluorescence. Consistent with the previous findings (5, 8, 44, 45) (46), the value for Mn^2+^ dissociation constant at the HAS in WT control Mn-depleted PSII complexes was shown to be at ~2.2 μM. Surprisingly, the K_m_ value for D1-E189K at physiological pH 6.0 appeared to be only slightly different from WT dissociation constant, ~2.9 μM indicating that Mn^2+^ binding at the HAS and consequent electron donation is not impaired in this mutant. If the affinity change at pH 6.0 would be the sole reason for D1-E189K inability to assemble the a large fraction of center we would expect the kinetics similar to D1-D170E mutant (**Fig. 2**), however it seems that slightly lower K_M_ at pH 6.0 could be only accounted for the decreased quantum efficiency of photoactivation during the early phase in D1-E189K mutant (**Fig. 2, Table 2**). On the opposite, Mn^2+^ affinity in D1-D170A (**Fig. S4, Table 4**), that has been shown to have a drastic effect on Mn^2+^ affinity at the HAS and fail to grow photoautotrophically (8), was previously estimated at ~50 μM Mn^2+^ possibly indicating on a dominant role of D1-D170 in the formation of the HAS in a comparison to D1-E189.

**Table 4.** Mn affinity at the HAS in purified Mn-depleted PSII from WT control, D1-E189K and D1-D170A. Obtained values were derived from the fluorescence plots fitted into Michaelis-Menten equation (**Fig. S4**) and based on Mn^2+^ electron donation capabilities in studied mutants. * Km value adapted from Nixon and Diner, 1992 (8)

| Strain | Mn<sup>2+</sup> affinity, $\mu M$ | |
| --- | --- | --- |
|  | pH 6.0 | pH 7.5 |
| WT control | 2.2 | 1.1 |
| D1-E189K | 2.9 | 0.9 |
| D1-D170A | 50* | 28.9 |

Lysin epossesses one of the highest side chain’s pKa values among all standard amino acids, for the sake of the current experiment the assay buffer pH was also set to pH 7.5. At elevated pH the affinity at the HAS in WT and D1-E189K was essentially the same again suggesting that D1-E189K is not involved in initial Mn^2+^ binding and oxidation at the HAS.

### 3.4. Accumulation of unstable photooxidant at the donor side of PSII

The observation that D1-E189K mutant has no small changes in Mn^2+^ affinity at the HAS (**Fig. S4, Table 4**) but yet reach only 40% of maximal O_2_ evolution activity after treatment with hydroxylamine (**Fig. 1B**) combined with the biphasicity of the assembly process (**Fig. 2**) may indicate on a ‘bifurcation’ of the photoactivation process with essentially two population of centers being formed, active and inactive. Previous analysis concluded that these mutations in this position result in cells containing both intact catalytic Mn clusters, plus significant fractions inactive centers containing photooxidizable Mn (20). To investigate if the mutations at the HAS indeed result in the formation of photooxidant at the donor site of apo-PSII (40, 41) we sought to partially photoactivate the cells to approximately 20-30% of maximal restoration level (**Fig. 2**) with 50 and 200 flashes for WT and D1-E189K respectively and estimate the stability of the photooxidant after the dark incubation. In the D1-A344Stop mutant, which fails to assemble active Mn_4_O_5_Ca centers, charge separation induces the formation of unstable photooxidized Mn^≥+3^ capable of serving as an oxidant of Q_A_^−^ and resulting in characteristic changes in Chl fluorescence relaxation kinetics with an increase in the proportion of fluorescence relaxation occurring more slowly (t_1/2_~ 240ms). Moreover, dark incubation resulted in the loss of non-functional Mn^≥+3^ (41).

Similarly, the purpose of the dark incubation (40 min) in the present experiments with partially photoactivated cells is to allow the re-reduction of unstable photooxidized Mn at the donor side, which might alter fluorescence relaxation kinetics (**Fig. S2, Table S2**). After the partial photoactivation WT control showed the reduction of the decay yield during the slow phase (55%) and appearance of the middle phase (1s, 27.4%) (**Fig. 4A**). The dark incubation of WT control cells has not significantly changed fluorescence relaxation kinetics indicating that the majority of photooxidized Mn at the donor side is assembled into functional and stable Mn cluster (**Fig. 4C**). Similarly to WT control, D1-E189Q mutant did not exhibit any significant changes in the fluorescence relaxation kinetics after the dark incubation (**Fig. S5 B&E, Table S4**).

**Figure 4.**
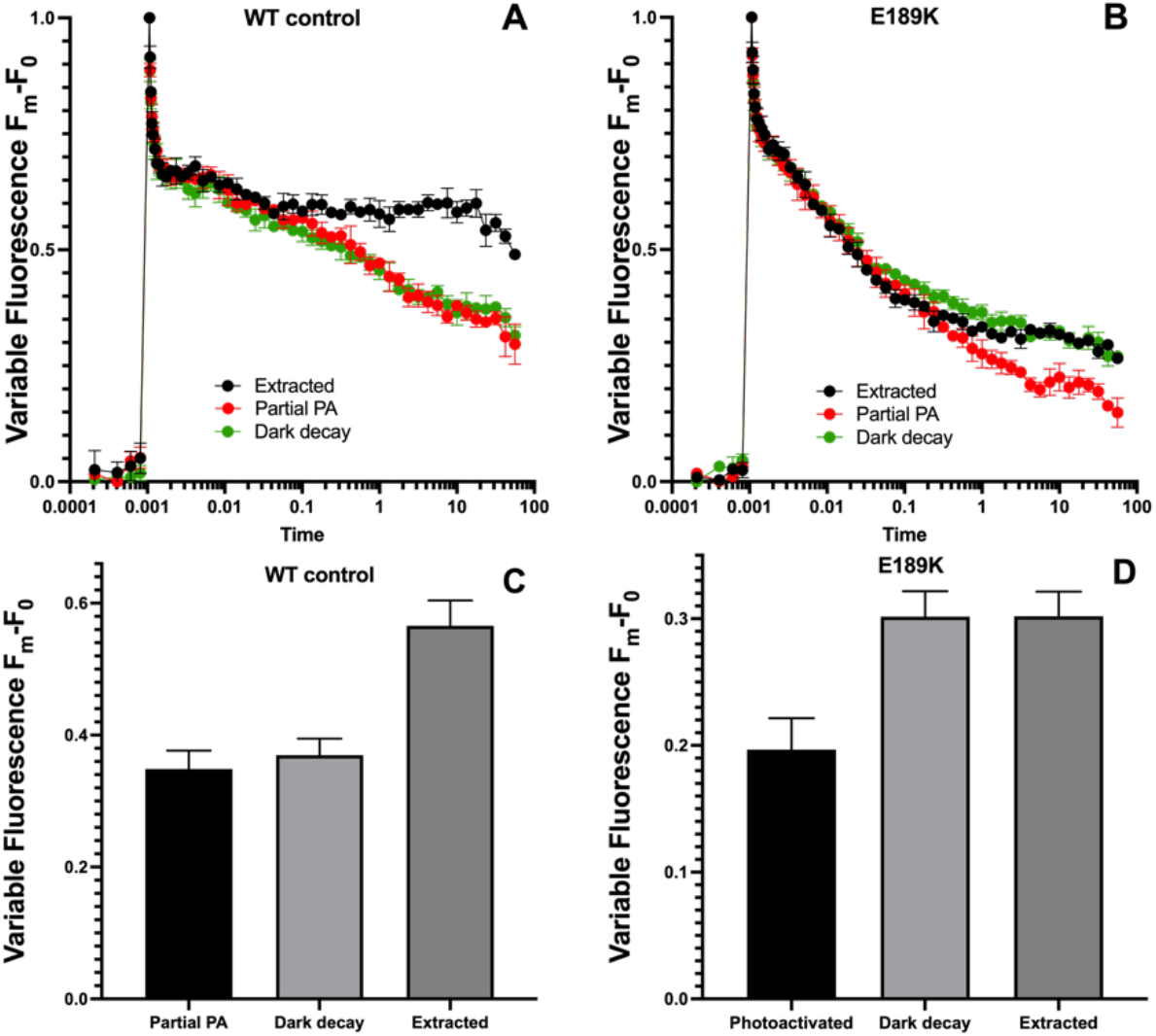
Q_A_^−^ reoxidation kinetics in partially photoactivated **(A)** WT control and **(B)** D1-E189K in the presence of DCMU. For the purpose of the partial photoactivation the sequence of 50 and 200 single turnover flashes was given to the HA-extracted WT control and D1-E189K cells respectively at uniform interval at 2Hz (500ms flash interval). Dark incubated sample was stored in complete darkness on a rotary shaker for 40 minutes prior the measurements. Panels **(C)** and **(D)** show the relative difference in the fluorescence relaxation kinetics in WT control and D1-E189K prior photoactivation, after photoactivation and after the dark incubation, the average of the last 8 fluorescence datapoints from panels **(A)** and **(B)** were taken to obtain the bar plots, higher level of fluorescence after the dark incubation indicates on a slower relaxation due to the absence of electron acceptor at the donor side of PSII. The measurements were performed after a single actinic flash of light. Samples were dark adapted for 5 minutes prior the measurements. Error bars represent SD with n ≥ 3

Photoactivation of D1-E189K (**Fig. 4B**) mutant with 200 flashes resulted in decreased time constant for the slow phase (100 s, 31%) compared to HA-extracted cells (>100 s, 41%) and increased half time of the middle phase from 144 ms and 18.5% to 457 ms and 27.6% of total yield respectively. Dark incubation caused the decay of unstable photooxidant at the donor side of D1-E189K mutant with consequent increase of the time constant and yield for the slow phase (>100 s, 44%) and decrease of the decay yield during the middle phase (16.7%) (**Fig. 4D**). Similarly, D1-D170A and D1-E189R mutants showed noticeable changes in the fluorescence relaxation after the dark incubation, indicating on a portion of unstable photooxidized Mn accumulated at the donor side of PSII, however to a much smaller degree compared to D1-E189K mutant (**Fig. S5, Table S4**).

## 4. Discussion

The primary event during light driven assembly of the Mn_4_O_5_Ca is the binding and photooxidation of a Mn^2+^ ion at the HAS of PSII (5, 6, 48). Previous mutagenesis studies (8, 24, 31, 35) identified D1-D170 as a key amino acid residue for Mn^2+^ binding and photooxidation, which represents the first step of photoactivation (**Scheme 1, A⟹B**). The crystal structure of holo-PSII reveals that the carboxyl group of D1-D170 provides one monodentate ligand to the ‘dangler’ Mn ion (Mn_4_) and one monodentate ligand the Ca^2+^ in the assembled complex. However, recent cryo-EM structures of apo-PSII show that this region of the WOC adopts a significantly different structural configuration compared to the fully assembled and active WOC (15, 16). Recently published apo-PSII structures identify electron density attributed to a heavy cation situated between the carboxylates of the D1-D170 and D1-E189, which was proposed to be an Mn^2+^ ion thus defining the HAS in apo-PSII. However, it is also possible that the observed density is a Ca^2+^, a reasonable conclusion given that D1-E189 ligates Mn1 and Ca^2+^ in an intact Mn_4_O_5_Ca cluster. The site directed mutants at D1-D170 showed a dramatic effect of these mutation on the ability to bind and oxidize Mn^2+^ by apo-PSII (7, 31), which ranges from complete loss of the function in D1-D170A to mutants capable of assembling up to 50% of WT O_2_ evolution activity in D1-D170E and D170H mutants (35). Less is known about the potential role of D1-E189 in forming the HAS, although early studies of D1-E189 mutants showed that many substitution mutants contain partially or misassembled Mn. Subsequent studies have shown that D1-E189 participates in a hydrogen bond network, modulates the redox potential of Y_Z_, and undergoes a conformational movement during the catalytic cycle proposed to allow substrate water delivery to the active site. Despite these extensive studies in the framework of water oxidation (20–23), until now, there has not been an attempt to evaluate the potential role of D1-E189 in the formation of the HAS and the assembly of the Mn_4_O_5_Ca.

### 4.1. Does D1-E189 to help form the high affinity site of PSII?

The electron donation experiment (**Fig. S4, Table 4**) in D1-E189K shows that this mutant possess the same Mn^2+^ affinity at the HAS indicating on a correct formation of photoactivation intermediates **A** and **B**, while D1-D170 **(Table 4)** mutations at the HAS inhibit initial Mn^2+^ photooxidation (35), thus preventing the electron donation to P_680_^+^ after the charge separation. Thus, analysis of the HAS characteristics of the D1-E189K, show that unlike mutations at D1-D170, the the Lys substitution of Glu189 does not affect the electron donation from Mn^2+^ towards Y_Z_ (**Fig. S4, Table 4**). Although a Lys substitution of D1-Asp170 has not been reported, the corresponding Arg mutation resulted in a five-fold reduction in affinity. Thus, we conclude that D1-E189 is not likely directly involved in forming the HAS. Consistent with previous results (20) D1-E189K mutation seems to have almost no effect on the O_2_ evolution rate, which reaches up to 80% of WT control rate (**Fig. 1**). While D1-E189Q mutation decreases the maximal rate to a larger extent, the net O_2_ rates are still relatively high and reach approximately 60% (20). Despite significantly high O_2_ rates in all studied D1-E189 mutants, an analysis of their photoactivation characteristics revealed severe defects. While D1-E189Q was able to restore initial oxygen evolution activity up to 70% after 15 minutes of continuous illumination, which is similar to the D1-D170E mutant. However, D1-D170E required twice as much time to reach same level of restoration (**Fig. 1B**). The photoactivation in the D1-E189K mutant was severely inhibited with the restoration level reaching 40% of initial O_2_ evolution activity respectively after 40 minutes of illumination. Unlike in the D1-D170E mutant where the ***Φ**_PA1_* was significantly lowered (**Table 2**), D1-E189K, D1-E189Q and D1-E189R (**Supplementary materials**) mutants re-assemble the Mn cluster at a much higher ***Φ**_PA1_* (**Fig. 2, Table 2**). However, after reaching 40% in D1-E189K and 80% in D1-E189Q the photoactivation almost stops and continues with a relatively low ***Φ**_PA2_*. Interestingly, D1-E189R mutant show very similar photoactivation properties to D1-E189K, suggesting the similar inhibitory role of positively charged amino acid on photoactivation (**Fig. S3**). The inability of the D1-E189 mutants to restore high level of WOC activity could indicate on the presence of internal damage or inhibition occurring either during the Mn-depletion procedure of photoactivation. Since all cyanobacterial strains were essentially treated the same and neither WT control nor D1-D170E strains showed similar assembly inhibition, it is unlikely that the treatment with 10mM of hydroxylamine caused the observed effect. Moreover, photoactivation was similarly inhibited during preliminary Mn-depletion experiments with 2mM of hydroxylamine in D1-E189K (**Fig. S6**). For all mutants the optimum interval between the photoactivation flashes yielding the maximum recovery of oxygen evolution by WOC was observed at 200-500 ms range (**Fig. S1, Table S1**), the flash interval experiments showed that the photoactivation intermediates in D1-E189K decay faster than in the wild-type. However, neither the overall low quantum efficiencies of photoactivation nor the increased rates of decay explain the inability of the D1-E189K to photoactivate such a large proportion of centers. It is worth mentioning that D1-E189K mutant is showing significant rates of O_2_ evolution prior the HA-extraction (**Fig. 1A**), however not able to restore this activity rapidly like WT control (**Fig. 1B**) while D1-E189Q similar to WT control is capable of rapid restoration of 80% of the net O_2_ rate (**Figs. 1B, 2**).

Analysis of the course of photoactivation under flash illumination revealed essentially two phases of the restoration curve. One phase, the main phase, corresponds to relatively high quantum efficiency of photoactivation, whereas the other phase involves a much lower quantum efficiency that, according to previous *in vitro* studies (37), can actually be a negative indicator of damage to the reaction centers during assembly. In the whole cell experiments conducted here, the main assembly phase in all mutants showed decreased quantum efficiency, ***Φ**_PA1_* (**Table 2**) suggesting that D1-E189 is important during initial steps of photoactivation. The reduced quantum efficiency, can be attributed to alterations in the HAS however, again these alterations does not explain the inability of D-E189 mutants to photoactivate with the constant rate similar to D1-D170E.

### 4.2. Does D1-E189 ligand participate in the binding of Ca^2+^?

D1-E189K mutant showed high quantum efficiency, ***Φ**_PA1_* early in the photoactivation that yields 40% of initial activity (**Fig. 2**) followed by the slow restoration phase, ***Φ**_PA2_* (**Table 2**). The reasons behind the fact that assembly significantly slows after a rapid phase of photoactivation, ***Φ**_PA1_* are intriguing and could be explained by the assumption that only a small fraction of the centers from the entire population of PSII are actually assembling functional WOC, while the remaining PSII centers are inhibited or severely slowed in assembly. This is consistent with the observation that unextracted D1-E189K cells exhibit significantly higher O_2_ evolution rate (80% of WT), indicating that a larger population of centers could assemble WOC over long period of time, which in turn indicates the reversible nature of observed inhibition. This and observation that D1-E189K can assemble a fraction of active WOC (**Figs. 2 and S1D**) at reasonably high quantum efficiency (**Table 2**) suggests that inability to assembly WOC by D1-E189K is not the result of a failure to photooxidize Mn^2+^ (**Fig. S4, Table 4**) or perform the dark rearrangement (Fig. **S1D**), but rather a result of inhibitory effect occuring over the course of illumination.

The change in the fluorescence relaxation kinetics in the presence of DCMU in Mn-depleted and partially photoactivated cells (**Fig. 4, S5**) indicate the accumulation of the photooxidant at the donor side of apo-PSII capable of charge recombination with Q_A_^−^. Considering the efficient photoactivation in WT and D1-E189Q cells (**Fig. 2**) there are no doubts that these strains accumulated functional WOC over the course of 50 and 100 flashes respectively. Although the charge recombination pathway also occurred after the partial photoactivation of D1-E189K and D1-D170E (**Fig. 4, S5**) mutants indicating the accumulation of the photooxidant, the nature of the photooxidant remains unclear. Consistent with altered quantum efficiency of photoactivation, ***Φ**_PA1_* (**Table 2**) due to the mutation at the HAS, aforementioned mutants could accumulate a population of oxidized carotenoids that slowly recombine with Q_A_^−^ (49) (**Fig. 4, S2, S5, Table 5**). However, this hypothesis again does not explain why D1-E189K mutant is not capable of acquiring a substantial fraction of active WOC during photoactivation (**Figs. 1B, 2**). Moreover, the retention of variable fluorescence indicates that this is not due to damage to the photochemical reaction center.

**Table 5.** Characteristics of chlorophyll fluorescence relaxation kinetics in partially photoactivated WT control and D1-E189K.

| Partially photoactivated cells in the presence of DCMU |  |  |  |  |  |
| --- | --- | --- | --- | --- | --- |
| Strain/State | | Fast phase: $\tau$/ms<br>(amplitude (%)) | Middle phase: $\tau$/ms<br>(amplitude (%)) | Slow phase: $\tau$/s<br>(amplitude (%)) | R |
| WT control | Extracted | 7.9 (14.5) | (0)- | > 100 (85.4) | 0.97 |
|  | Partial Photoactivation | 8.5 (17.7) | 1050 (27.4) | > 100 (54.9) | 0.99 |
|  | Dark incubated | 8.3 (19.9) | 918 (22.3) | > 100 (57.9) | 0.98 |
| D1-E189K | Extracted | 5.9 (40.5) | 144 (18.5) | > 100 (40.9) | 0.99 |
|  | Partial Photoactivation | 7.4 (41.2) | 457 (27.6) | 100 (31.1) | 0.99 |
|  | Dark incubated | 7.9 (39.2) | 440 (16.7) | > 100 (44.1) | 0.99 |

Alternatively, the photooxidant accumulated at the donor side of apo-PSII could be attributed to mis-assembled photooxidized Mn as predicted earlier by Chu *et. al* (10, 24), Hwang *et*. al (35) and observed by Cser *et. al* in D1-A344stop mutant (35, 41). Considering that the quantum efficiency of photoactivation, ***Φ**_PA1_* in D1-E189K mutant in fact was less severely affected compared to D1-D170E mutant and no significant effect on Mn^2+^ affinity at the HAS in D1-E189K was observed (**Fig. S4, Table 4**), we tentatively assign the accumulated photooxidant at the donor side of apo-PSII to misassembled non-functional photooxidized Mn. The assumption that D1-E189K mutant accumulate non-functional Mn at the donor side in the majority of centers would explain the inability of this mutant to restore a large fraction of active WOC during relatively short course of photoactivation (**Figs. 1B and 2**). Although the mechanism of inappropriate Mn accumulation is unclear, one hypothesis could be derived from the fact that besides Mn1, D1-E189 coordinates Ca^2+^ in intact WOC (14). Considering the general agreement that Ca^2+^ is crucial for photoactivation (12, 18, 37, 50) the actual mechanism how Ca^2+^ facilitates the photoassembly remains elusive. Binding of Mn^2+^ to the HAS with simultaneous binding of Ca^2+^ to an adjacent binding site facilitates the formation of the [Mn^2+^-(OH)-Ca^2+^] complex by inducing deprotonation of a water ligand of Mn^2+^. Calcium lowers the pK_a_ for water ligand, which is controlled by a nearby base B^−^ that serves as a primary proton acceptor with a p*K*_a_ dependent on Ca^2+^ bound to its effector site (51). It is possible that D1-E189 ligand supports Ca^2+^ binding at its effector site in apo-PSII during early photoactivation steps. Considering the strong competition between Mn^2+^ and Ca^2+^ for the CES (37, 38) shown during *in vitro* experiments, and the observed negative effect of D1-E189 mutations on the photoactivation it is reasonable to assume that D1-E189K and potentially D1-E189R (**Fig. S3**) mutations could alter the binding of Ca^2+^ to the CES, thus even more increasing the Mn^2+^ inhibitory effect (37, 38). According to Chen *et al*. (18), Ca^2+^ prevents inappropriate binding of high valency non-functional Mn ions at the donor side of apo-PSII during the photoactivation, an effect that can also occur at high light intensities (43). Interestingly, partially assembled Mn clusters were inferred by Cser and these appeared to be unstable in the dark, which resulted in the loss of charge recombination pathways in D1-A344stop mutant (41). Consistent with earlier observations, D1-E189K showed noticeable change in the fluorescence decay kinetics after partially photoactivated samples were incubated in the dark (**Figs. 4B & D**), while WT (**Figs. 4A & C)** and D1-E189Q (**Fig. S5**) strains have not shown any changes in fluorescence relaxation kinetics.

Assuming that in D1-E189K mutant only a fraction of the centers successfully ligated Ca^2+^ prior the photoactivation, the entire population of centers undergo essentially two different pathways of photoactivation. The bifurcation of the photoactivation process represent the ratio distribution of assembled centers from the common pool of inactivated centers by hydroxylamine treatment and could be considered a competition between productive photoactivation, when Ca^2+^ is bound at the CES, and nonproductive photoactivation in the absence of Ca^2+^. The results observed during flash number experiments in D1-E189K mutant indicate the ratio of non-functional Mn is much higher compared to WT control and reaches up 60%, however D1-E189Q assembled approximately 80% of the centers correctly and only smaller fraction of centers were inactivated (**Fig. 2**). The similarity in rate of the dark rearrangement, *K_A_* (**Table S1**), among the mutants suggests that the assembly of appropriate Mn cluster occurs with similar kinetics, which in principle is not much different from WT control. The main difference between these mutants is the distribution of the assembly pathways yielding different amounts of functional and nonfunctional centers. Previous findings indicate that inappropriate Mn does not cause permanent damage to the PSII centers and can be re-reduced by HA treatment allowing further photoassembly (35). Our results show that D1-E189K reassemble non-functional Mn although at the slow rate (**Table 2**); that would explain its ability to show high O_2_ evolution rates prior the HA treatment.

### 4.3. Does the accumulation of non-functional Mn in PSII irreversibly inhibit photoactivation?

As mentioned earlier the unique property of D1-E189K and D1-E189Q is a significant rate of oxygen evolution among all PSII mutants. In principle this feature indicates that the cells are capable of coping with the inhibition caused by the formation of non-functional Mn structures (35, 43) as discussed earlier. However, it is not clear if non-functional Mn could be removed from the apo-PSII without the initiation of PSII repair process, which involves the disassembly of PSII monomer and replacement of the D1 protein (52–54). The evidence that the PSII centers populated with non-functional Mn at the WOC site could be restored by hydroxylamine treatment (35) and by essentially the same electron donation capability in D1-E189K (**Fig. S4, Table 4**) imply that the oxidative state of inhibitory Mn at the WOC most likely higher than Mn^2+^ and that the reduction of such Mn results in its dissociation from the inhibition site. Interestingly, in the absence of the PsbO protein, Mn_4_O_5_Ca cluster tends to be reduced in the dark, which could result in complete loss of photosynthetic function in this mutant (42, 55). The fact that the ΔPsbO mutant is capable of significant O_2_ evolution suggests that the process of cluster reduction and re-assembly happens concurrently under illumination. In principle, **Φ***_PA2_* in D1-E189K and to a large extent in D1-E189R (**Fig. S3**) mutant could indicate the reduction of ‘wrong’ Mn followed by new assembly trial over the course of 6400 photoactivation flashes (**Fig. 2**). Since the Mn cluster in ΔPsbO can be reduced by endogenous reductants in the lumen, non-functional Mn formed during the assembly in D1-E189K mutant could potentially decay in the dark. Additional evidence for the reduction of partially assembled Mn cluster in the dark come from D1-A344Stop mutant, where 40-minute dark incubation resulted in significant reduction of the charge recombination between Q_A_^−^ and photooxidized Mn at the donor side of PSII (41). Consistent with these observations, the changes in fluorescence relaxation kinetics were not observed after the dark incubation in WT control (**Fig. 4A & C**). The consistency in fluorescence decay kinetics prior to and after the dark incubation indicate that the majority of assembled centers over the course of 50 flashes were formed in fully functional and stable Mn clusters.

Among the studied strains only D1-E189Q mutant showed similar photoactivation to WT control (**Figs. 2, S1**) and comparable Q_A_^−^ reoxidation kinetics in the samples prior HA treatment and after Mn extraction (**Fig. S2**). Similar to WT control, D1-E189Q exhibited virtually no difference in fluorescence decay kinetics prior the dark incubation and after **(Figs. S4B & E)**. Despite the difference in p*K_A_* between Glu and Gln, D1-E189Q seems to be capable of supporting correct assembly of the Mn cluster similar to WT control. Unlike in WT control and D1-E189Q strains **(Figs. 4, S5)** D1-E189K (**Fig. 4**) and D1-D170E **(Figs. S5)** accumulated a portion of unstable photooxidant, that we assign to non-functional Mn, at the donor site after the partial photoactivation. Similar to what has been observed in D1-A344Stop prior and after the dark incubation, D1-D170E and D1-E189K mutants showed reduction in fluorescence decay rate after 40-minute dark incubation (**Fig. S5, 4**). The slowdown in the fluorescence decay after the incubation in the dark suggests that these mutants lost a fraction of redox active Mn involved in the charge recombination between Q_A_^−^ and the donor side of PSII. These observations support previous hypothesis that during the photoactivation in D1-E189K mutant the assembly of the Mn cluster yields two types of centers, active centers capable of water oxidation and non-functional but yet redox active centers containing at least a single Mn^3+^ ion at the donor site of PSII (24). Importantly, the accumulation of the non-functional Mn by D1-E189 mutants while retaining the fluorescence relaxation kinetics after the reduction of the non-functional Mn indeed indicate on the absence of irreversible damage to the reaction centers similar to D1-D170V mutant (35). The ability of this mutants to assemble up to 80% of active centers in regard to WT control and the evidence that inappropriate Mn could be removed from the centers we suggest that the formation of non-functional Mn clusters in D1-D170E and D1-E189K mutants is reversible and in a long run does not prevent these mutants from assembling a significant portion of active PSII. However, verification of the tentatively assigned “inappropriate Mn” (18), will require further *in vitro* studies with purified PSII core complexes for careful investigation of the phenomenon.

## 5. Conclusions and model

While mutation of the D1-E189 ligand has a strong impact upon photoactivation, it is not disrupt the HAS in assays monitoring Mn^2+^ binding and oxidation. Unlike most D1-D170 mutants (8), most of the D1-E189 mutants are capable in accumulation of a large fraction (≥50%) of PSII centers containing photooxidizable Mn ions, although many of these are not capable of oxygen evolution activity (20). Given that D1-E189K has a very similar Mn^2+^ affinity at the HAS, but is highly impaired in photoactivation, we conclude that defects in the assembly of Mn_4_O_5_Ca cluster in D1-E189K occur after the initial Mn^2+^ oxidation. Given its role in binding Ca^2+^ in the fully assembled cluster and the propensity of D1-E189 mutants to form inactive clusters due to Mn oxide formation, we conclude the mutational defect in assembly is due to the perturbation of Ca^2+^ function during photoactivation. The observation of a cation situated between D1-Asp170 and D1-E189 in the apo-structure (16), it is likely that Ca^2+^ occupies this position, which constitutes the CES. It is worth noting that lysine substitutions for carboxylates in the Ca^2+^-binding EF hand structure do not abolish Ca^2+^ binding, although the affinity is diminished (56). According to the model shown in **Fig. 5**, the D1-E189K mutation perturbs the Ca^2+^-mediated formation of intermediate ‘**C**’ and/or the quantum efficiency of the photoconversion of **C⟹D** that leads to the formation of a di-μ-oxo bridged binuclear Mn^3+^, proposed to be the first stable photoactivation intermediate, (“**D**”). Previous analysis has shown that Ca^2+^ indeed stabilizes the intermediates during the dark rearrangement (19). Recent computational results (17, 57, 58) are consistent with x-ray crystallographic results (59) indicating that positions Mn1 and Mn2 in the assembled complex bind for Mn^3+^ more strongly than does the HAS. It is proposed that the rearrangement involves ion translocations to these more stable sites (59, 60). In this context, Ca^2+^ at the CES, facilitates the translocational rearrangement of the [Mn^3+^-(OH)-Ca^2+^] species (61) positioning the Mn^3+^ ion in a thermodynamically more stable site corresponding to the Mn(1) and Mn(2) locations in the intact PSII. This conformational folding produces a configuration resembling the fully assembled configuration probably similar to the configuration produce by Mn photooxidation in PSII crystals that had been extracted and re-illuminated (59). The role of Ca^2+^ is two-fold: 1) blocking the binding and photooxidation of Mn^2+^ at the CES which would otherwise to inactive Mn oxide formation and 2) tethering and thereby guiding the relocation of Mn^3+^ produced at the HAS into the correct site. The relocation would open the HAS for the second Mn^2+^ binding and photooxidation and a similar repositioning of this second ion as it joins the first to form the stable binuclear intermediate located at Mn(1) and Mn(2) and accompanied by protein structural rearrangements involving the CP43 e-loop and the D1 c-terminus.

**Figure 5.**
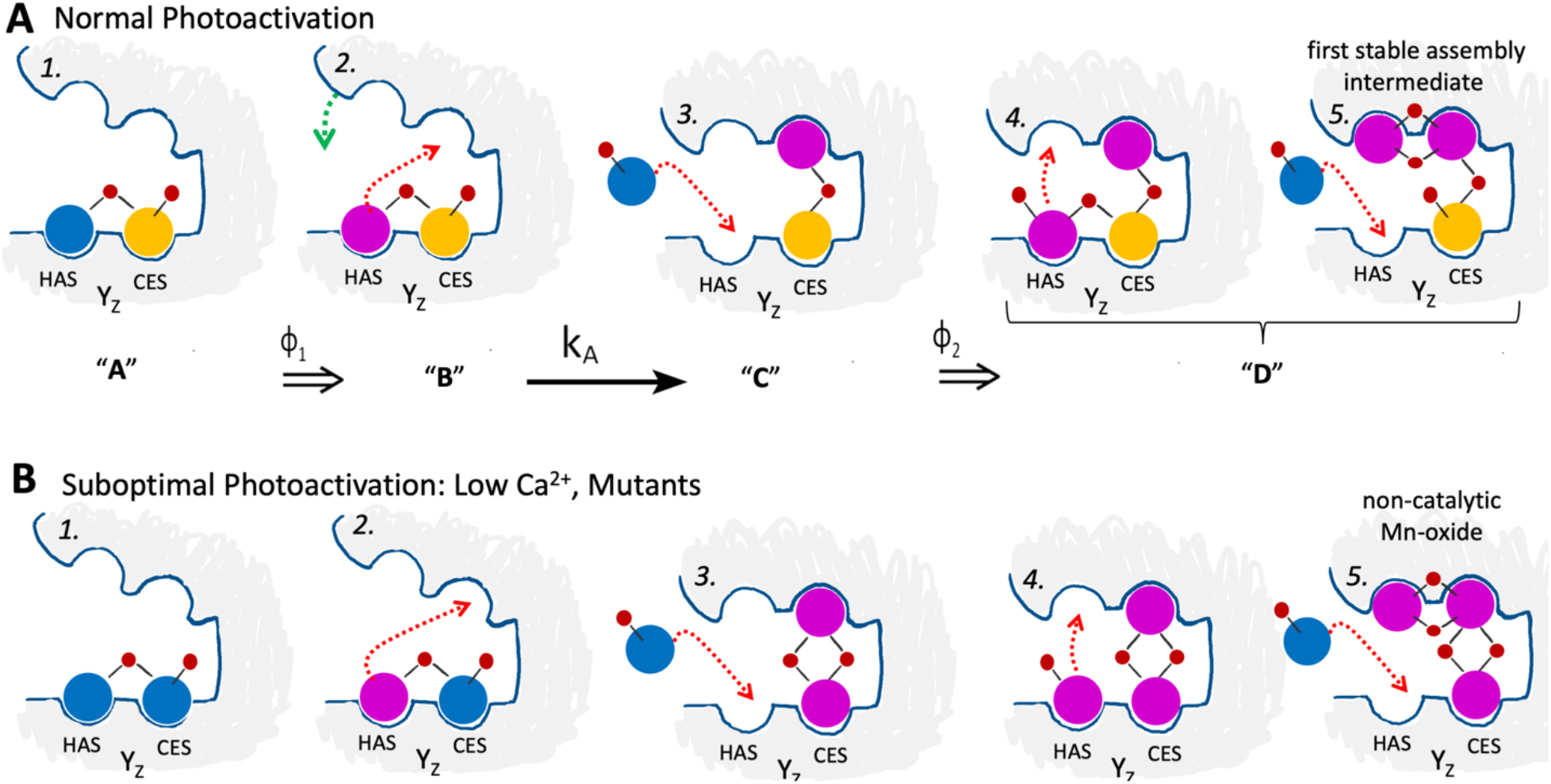
Models of normal photoactivation versus mis-assembly of Mn oxides under suboptimal conditions. (**A**) Initial steps of the photoassembly at optimal conditions. ***(1.)*** Formation of intermediate “**A**” **(Scheme 1)** where Ca^2+^ (yellow circle) bound to the Ca^2+^ effector site (CES) and Mn^2+^ (blue circle). In this moel, Ca^2+^ normally functions to prevent inappropriate Mn^2+^ as well as guiding the incoming Mn^2+^ via (hydr)oxo bridges (smaller red circles) derived from the hydration sphere of incoming cations. The Mn^2+^ is bound to the high affinity site (HAS) forming [Mn^2+^- (OH)-Ca^2+^] intermediate, ***(2.)*** formation of the intermediate “**B**”, first photooxidized Mn^3+^ (pink circle) undergoes a cooperative ion translocation (red dashed line) and conformational changes (green dashed line) (**B→C**) forming ***(3.)*** intermediate “**C**” vacating the HAS allowing binding of new Mn^2+^ for further photooxidation ***(4.)***. After the second photooxidation the Mn^3+^ binuclear intermediate “**D**” has been formed *(5)* allowing further Mn_4_CaO_5_ assembly. (**B**) Suboptimal photoactivation where Ca^2+^ binding is affected by low concentration (37) or perturbed Ca^2+^ binding site by mutation (e.g. D1-E189K). (***1***.) Both CES and HAS are each occupied by Mn^2+^. In this case, charge separation (***2***.) oxidizes Mn^2+^ at the HAS, and further charge separation (***3***.) oxidizes Mn^2+^ bound at the CES. The CES is located near Y_z_ suggesting that photooxidation of Mn^2+^ at the CES could even occur first (***4***.). Binding and photooxidation of additional Mn^2+^ results in multinuclear yet non-functional Mn^3+/4+^ cluster (*5*). Current models suggests both processes occurring in both WT control and D1-E189K mutants but with different ratio distribution. In WT control the normal photoactivation (**A**) is predominant with the lower ratio of suboptimal photoactivation (**B**) resulting in high photoactivation yield (**Figs. 1 and 2**). In D1-E189K the suboptimal photoactivation (**B**) is predominant due to the perturbed CES resulting in lower photoactivation yield, but the slow reversibility of Mn oxide inactivation by reduction *in vivo* allows assembly of the majority of active centers over the long period of time as observed in our experiments.

Photoactivation under suboptimal Ca^2+^ concentrations or in mutants with a perturbed CES lead inactivation due to “inappropriate Mn” (18). Partial impairment of Ca^2+^-function as with the D1-E189K substitution leads bifurcation of the photoactivation process resulting in essentially two populations of PSII centers: one where Ca^2+^ was successfully mediated Mn^3+^ translocation to form and active WOC and the other where Mn^2+^ bound to the CES and led to the formation of non-catalytic Mn-oxide formation (43, 62).

## Abbreviations

HAS: high affinity site
HA: hydroxylamine
P680: primary electron donor of the reaction center
PSII: Photosystem II
WOC: H_2_O-oxidizing complex
Y_z_: redox-active tyrosine secondary electron donor serving as reactive interface between the primary donor and the WOC.

## Supplementary results

### 5.1 Dark molecular rearrangement and the stability of photoactivation intermediates

According to the “two-quantum model” of photoactivation (**Scheme 1**), the formation of the first stable intermediate (intermediate **D**) involves two photoacts separated by a light-independent molecular rearrangement (4, 36). This rate-limiting rearrangement, combined with the limited stability of the photoactivation intermediates, results in a bell-shaped curve when the formation of active PSII is promoted by a fixed number of flashes given at different flash intervals, measuring the extent of recovery of O_2_ evolution after the flash sequence (34, 35, 37, 39, 63).

For the present experiments, a fixed number of flashes, was determined for each strain to give approximately 50% recovery at optimal flash spacing. This number of flashes was given to HA-extracted samples at intervals ranging from 50 millisecond to 10 seconds for each of the strains and the plotted yields at each flash frequency produced the ‘bell-shaped’ curves shown in Figure 3. To estimate time constants for the dark rearrangement, *k_A_*, and the decay of labile intermediates, *k_D_*, the flash interval photoactivation data was fit into the equation derived from the two-quantum model (35, 37, 39). The values for the dark rearrangement and the decay of intermediates in terms of half-times for WT control were 66 ms and 1.8 s respectively (**Table 3**). All mutants studied showed similar moderately slowed values for the dark rearrangement, *k_A_* ranging from 78–82 ms, the rate-limiting, molecular rearrangement (**Scheme 1, B→C**). Recent evidence suggests that the dark rearrangement, *k*_A_, involves a conformational change of the metal-binding polypeptide structure of the WOC (15, 37, 64). Since all mutants exhibit similarly decreased *k*_A_ value compared to WT control, yet had markedly different extents of assembly, the reason behind the inability of D1-E189K to assemble a large fraction of the PSII centers (**Figs. 1B and 2D**) is not associated with a slowed the rate of the molecular rearrangement (38, 42, 65, 66). Interestingly, all mutants exhibited decreased stability of the labile photoactivation intermediates (**Fig. 3, S1B and Table 3**). This result is consistent with D1-E189 providing as stabilizing interaction, either direct or indirect, with the Mn^3+^ ion produced during the initial photooxidation at the HAS **(A⟹B)**.

**Figure S1.**
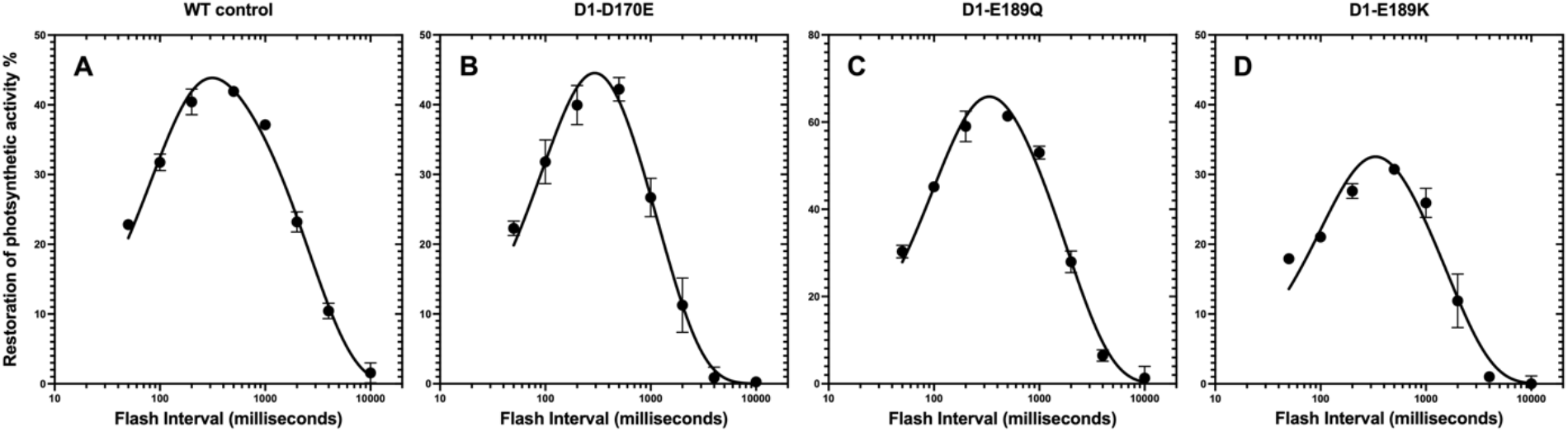
Photoactivation of HA-extracted cells as a function of flash interval. Flashes were given at six different equal flash intervals to HA-extracted WT control (Panel A), D1-D170E (Panel B), D1-E189Q (Panel C), D1-E189K (Panel D). The curves represent fits to Eq. 2 derived from the two-quantum model shown in Scheme 1 (39). The number of photoactivation flashes was determined based on the flash number experiment (Fig. 2) to produce approximately 50% restoration of activity. This corresponded to 150, 250, 350 and 500 for WT control, D1-E189Q, D1-E189K, and D1-D170E respectively. The percentage of O_2_ evolution recovery was calculated as a fraction from the O_2_ evolution rate prior HA extraction which were 600 ± 20, 369 ± 27, 520 ± 32 and 402 ± 12 of O_2_ (mg Chl)^−1^ h^−1^, for WT control, D1-D170E, D1-E189Q and D1-E189K respectively. Data were fit to Eq. 1 for parameter estimation. Error bars represent SD with n ≥ 3.

**Table S1.**
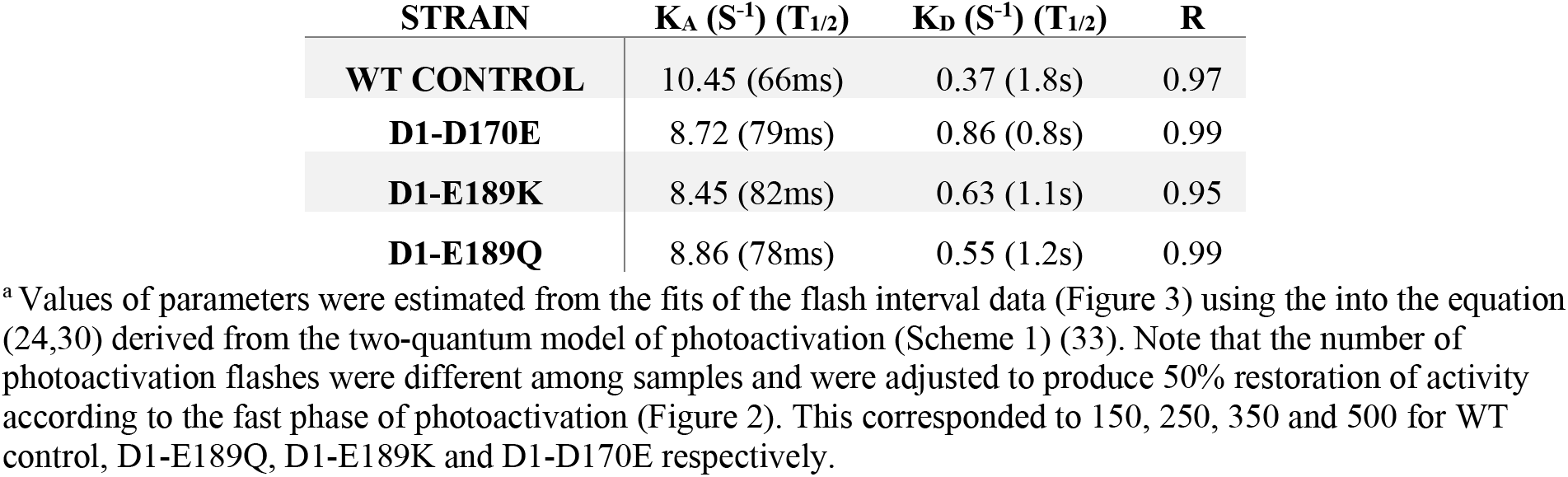
Dark rearrangement, kA and decay of intermediates, kD, parametersof WT control, D1-D170E, D1-E189K, D1-E189R, and D1-E189Q mutant strains.

| STRAIN | $K_A$ (S<sup>-1</sup>) (T<sub>1/2</sub>) | $K_D$ (S<sup>-1</sup>) (T<sub>1/2</sub>) | R |
| --- | --- | --- | --- |
| WT CONTROL | 10.45 (66ms) | 0.37 (1.8s) | 0.97 |
| **D1-D170E** | 8.72 (79ms) | 0.86 (0.8s) | 0.99 |
| **D1-E189K** | 8.45 (82ms) | 0.63 (1.1s) | 0.95 |
| **D1-E189Q** | 8.86 (78ms) | 0.55 (1.2s) | 0.99 |

<sup>a</sup> Values of parameters were estimated from the fits of the flash interval data (Figure 3) using the into the equation (24,30) derived from the two-quantum model of photoactivation (Scheme 1) (33). Note that the number of photoactivation flashes were different among samples and were adjusted to produce 50% restoration of activity according to the fast phase of photoactivation (Figure 2). This corresponded to 150, 250, 350 and 500 for WT control, D1-E189Q, D1-E189K and D1-D170E respectively.

### 5.2 Fluorescence relaxation kinetics

The acceptor side forward electron flow from Q_A_^−^ to Q_B_ site does not seem to be affected by Mn-depletion nor after photoactivation, even in the mutants (**Figs, S2A and S2E**) indicating an absence of acceptor side damage. However, the kinetics of charge recombination between the donor and acceptor side observed in the presence of DCMU in HA-treated cells indicate a significant alterations at the HAS in studied mutants. The D1-D170E and D1-E189K mutants displayed nearly complete decay of Q_A_^−^ within 1s following HA-extraction of the Mn_4_CaO_5_ (**Fig. 2F, Table S2**). In contrast, the decay of fluorescence WT control was not complete even after over 100s. The longer lifetime of Q_A_^−^ in the WT is due to the presence of Mn^2+^ at the HAS at the moment of the charge separation permitting efficient reduction of P_680_^+^ and thereby trapping the center in the high fluorescent Y_Z_P_680_Q_A_^−^ state (5, 20, 67). Although the Mn_4_CaO_5_ has been removed in the samples, there is a sufficient concentration of Mn^2+^ in the cells to allow full reassembly and therefore it is reasonable that Mn^2+^ can occupy the HAS in these samples.

**Figure S2.**
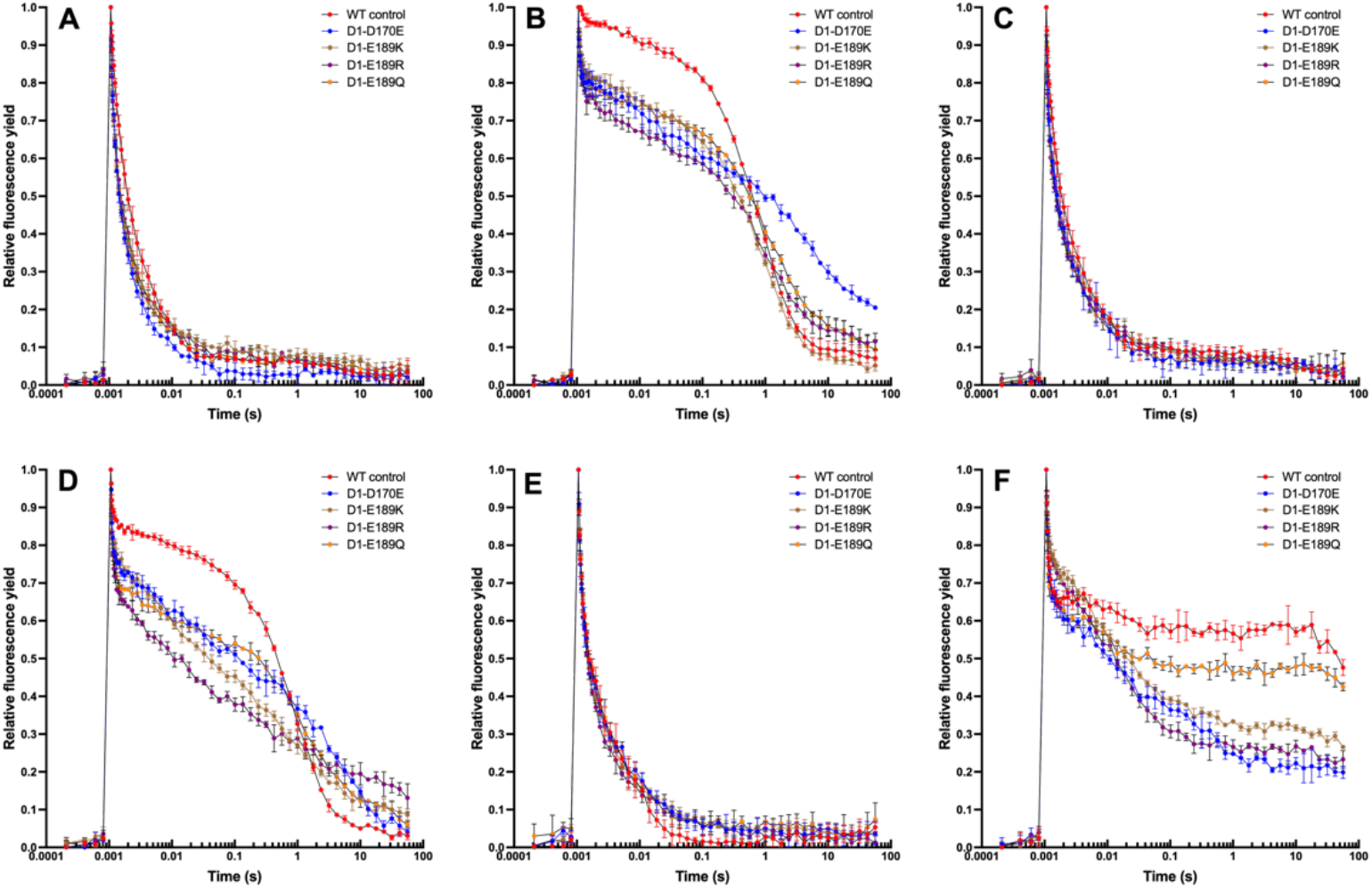
Q_A_^−^ reoxidation kinetics in WT control, D1-D170E, D1-E189K, D1-E189R, and D1-E189Q strains in the (**A**) absence of DCMU, (**B**) presence of DCMU, (**C**) re-photoactivated cells in the absence of DCMU, (**D**) re-photoactivated cells in the presence of DCMU, **(E)** Mn-depleted cells in the absence of DCMU, **(F)** Mn-depleted cells in the absence of DCMU. For the purpose of the photoactivation (**C and D**) the sequence of 2000 single turnover flashes was given to the HA-extracted cyanobacterial cells at uniform interval at 2Hz (500ms flash interval). The measurements were performed after a single actinic flash of light. Samples were dark adapted for 5 minutes prior the measurements. Error bars represent SD with n ≥ 3.

**Table S2.** Characteristics of chlorophyll fluorescence relaxation kinetics in untreated and re-photoactivated WT control, D1-D170E, D1-E189K, D1-E189R and D1-E189Q cells.

| Strain | Fast phase: $\tau$/ms<br>(amplitude (%)) | Middle phase: $\tau$/ms<br>(amplitude (%)) | Slow phase: $\tau$/s<br>(amplitude (%)) | R |
| --- | --- | --- | --- | --- |
| Intact Mn<sub>4</sub>O<sub>5</sub>Ca cluster in the absence of DCMU |  |  |  |  |
| WT control | 0.4 (86.9) | 4.29 (11.1) | 8.3 (1.9) | 0.99 |
| D1-D170E | 0.3 (96) | 7.4 (3.5) | 86.6 (0.5) | 0.99 |
| D1-E189K | 0.2 (97.3) | 5.2 (2) | 34.8 (0.6) | 0.99 |
| D1-E189R | 0.3 (95.8) | 4.9 (3.3) | 6.9 (0.9) | 0.98 |
| D1-E189Q | 0.26 (96.8) | 5.9 (2.6) | 36.5 (0.7) | 0.98 |
| Intact Mn<sub>4</sub>O<sub>5</sub>Ca cluster in the presence of DCMU |  |  |  |  |
| WT control | 1.6 (11) | (0)- | 0.75 (89) | 0.99 |
| D1-D170E | 13.9 (24.8) | 3008 (47.9) | >100 (27.2) | 0.99 |
| D1-E189K | 3 (16) | (0)- | 0.8 (75) | 0.98 |
| D1-E189R | 3.9 (25.4) | (0)- | 1.6 (74.5) | 0.98 |
| D1-E189Q | 0.93 (28.9) | (0)- | 1 (71) | 0.99 |
| Re-photoactivated cells in the presence of DCMU |  |  |  |  |
| WT control | 0.8 (21.8) | (0)- | 0.7 (78.2) | 0.99 |
| D1-D170E | 0.9 (23.5) | 308 (26.9) | 4.7 (49.6) | 0.99 |
| D1-E189K | 5.6 (37.5) | 627 (49.4) | >100 (13.1) | 0.99 |
| D1-E189R | 3.7 (38.1) | 269 (33.7) | 96.9 (28.1) | 0.99 |
| D1-E189Q | 2.36 (24) | 1011 (63.3) | 89.6 (12.6) | 0.99 |
| Mn-depleted cells in the presence of DCMU |  |  |  |  |
| WT control | 7.9 (14.5) | (0)- | > 100 (85.4) | 0.99 |
| D1-D170E | 6.5 (39.5) | 446 (27.9) | > 100 (32.5) | 0.99 |
| D1-E189K | 5.9 (40.5) | 144 (18.5) | > 100 (40.9) | 0.99 |
| D1-E189Q | 0.27 (72.8) | 15.8 (6.5) | > 100 (20.7) | 0.98 |

### Supplementary figures

**Figure S3.**
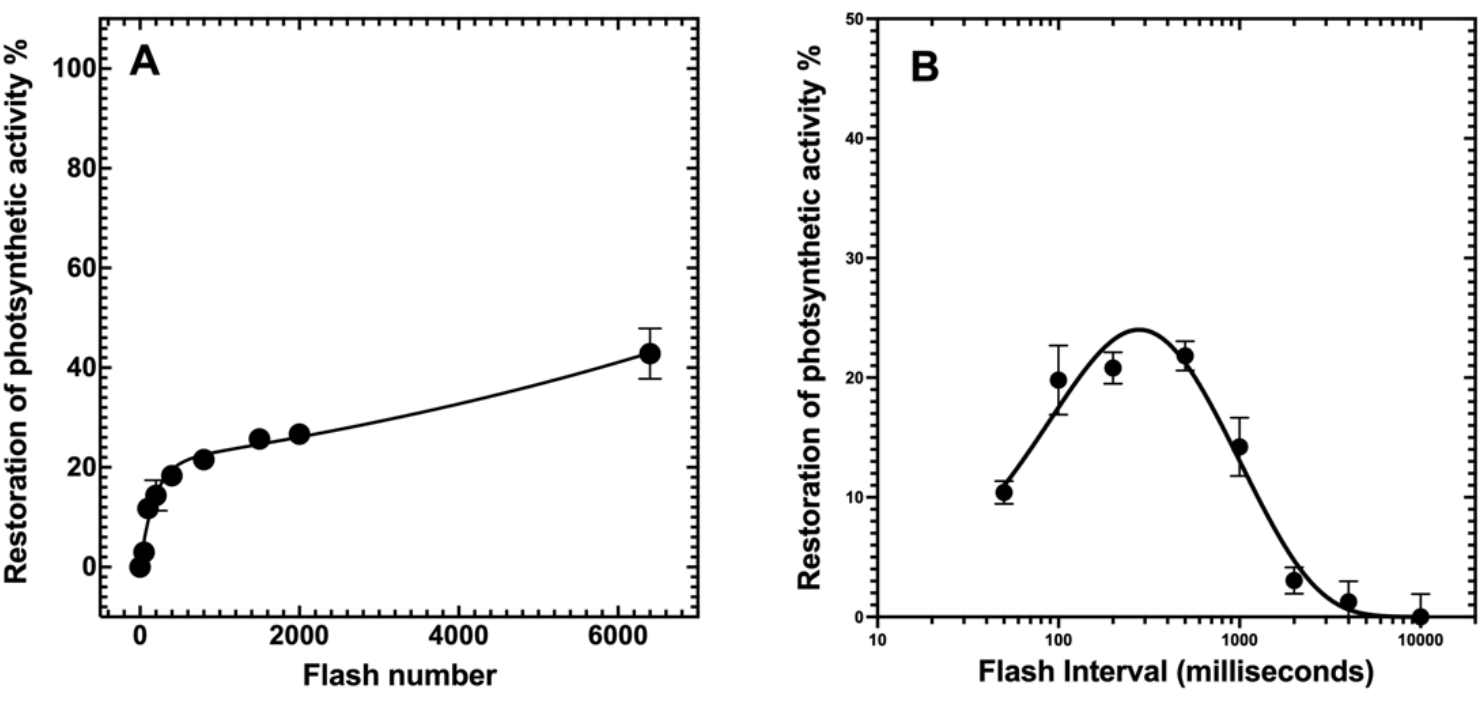
Photoactivation of HA-extracted cells as a function of (**A**) single-turnover flash number. Series of xenon lamp flashes were given at uniform frequency of 2 Hz (500ms flash interval) to hydroxylamine-treated D1-E189R cells. Kinetic fit parameters of ***Φ**_PA1_* = 5.7 × 10^−3^ *and **Φ**_PA2_* = 1.1 × 10^−4^ obtained from the data from the was fit into Equation 1. (**B)** Photoactivation of HA-extracted cells as a function of flash interval. Flashes were given at six different equal flash intervals to hydroxylamine-treated D1-E189R cells. The number of photoactivation flashes was determined based on the flash number experiment (**Fig. S1A**) to produce approximately 50% restoration of activity and corresponded to 350 flashes. Data were fit to Eq. 2 for parameter estimation. The percentage of O_2_ evolution recovery was calculated as a fraction from the O_2_ evolution rate prior HA extraction equals 262 ± 9 μmol of O_2_ (mg Chl)^−1^ h^−1^. Error bars represent SD with n ≥ 3

**Figure S4.**
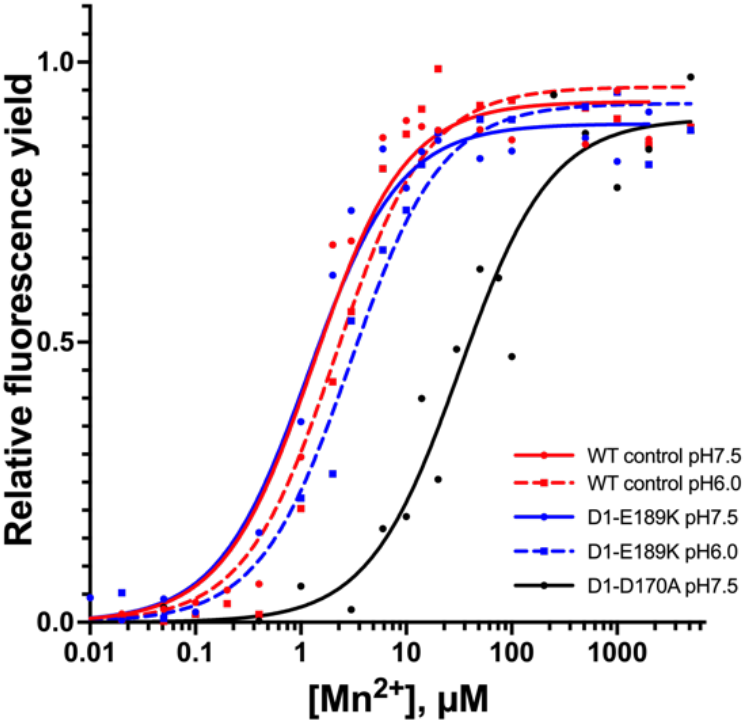
Electron donation in purified Mn-depleted PSII as a function of Mn^2+^ concentration. The variable fluorescence emitted by PSII from WT control, D1-E189K and D1-D170A strains at pH6.0 (dashed line) and pH7.5 (solid line) was measured after the excitation with saturating laser flashes (5ns, 300ms apart) (43). Sample contained 5μg of Chl\mL, 10μM 2,6-DCBQ as an external electron acceptor, pH 6.0). Fluorescence data was normalized to 1 and fit into Michaelis-Menten equation to derive K_m_ values. Every single data point represents an average of 3-5 technical repeats

**Figure S5.**
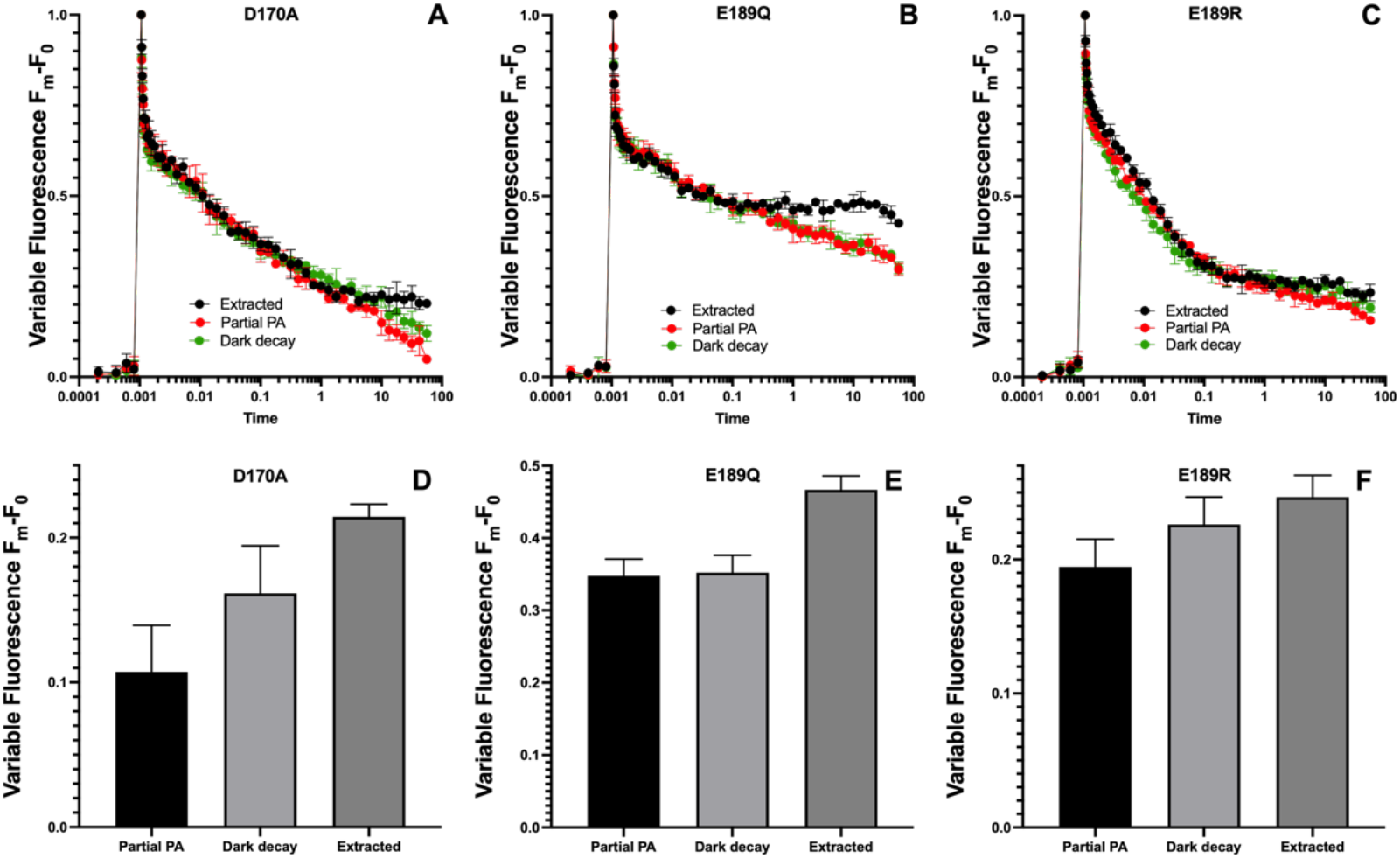
Q_A_^−^ reoxidation kinetics in partially photoactivated **(A)** D1-D170E, **(B)** D1-E189Q, and **(C)** D1-E189R mutants in the presence of DCMU. For the purpose of the partial photoactivation the sequence of 300, 200 and 200 single turnover flashes was given to the HA-extracted cells at uniform interval at 2Hz (500ms flash interval), respectively. Dark incubated sample was stored in complete darkness on a rotary shaker for 40 minutes prior the measurements. Panels **(D)**, **(E)** and **(F)** show the relative difference in the fluorescence relaxation kinetics in WT control and D1-E189K prior photoactivation, after photoactivation and after the dark incubation, the average of the last 8 fluorescence datapoints from panels **(A), (B)** and **(C)** were taken to obtain the bar plots, higher level of fluorescence after the dark incubation indicates on a slower relaxation due to the absence of electron acceptor at the donor side of PSII. The measurements were performed after a single actinic flash of light. Samples were dark adapted for 5 minutes prior the measurements. Error bars represent SD with n ≥ 3

**Figure S6.**
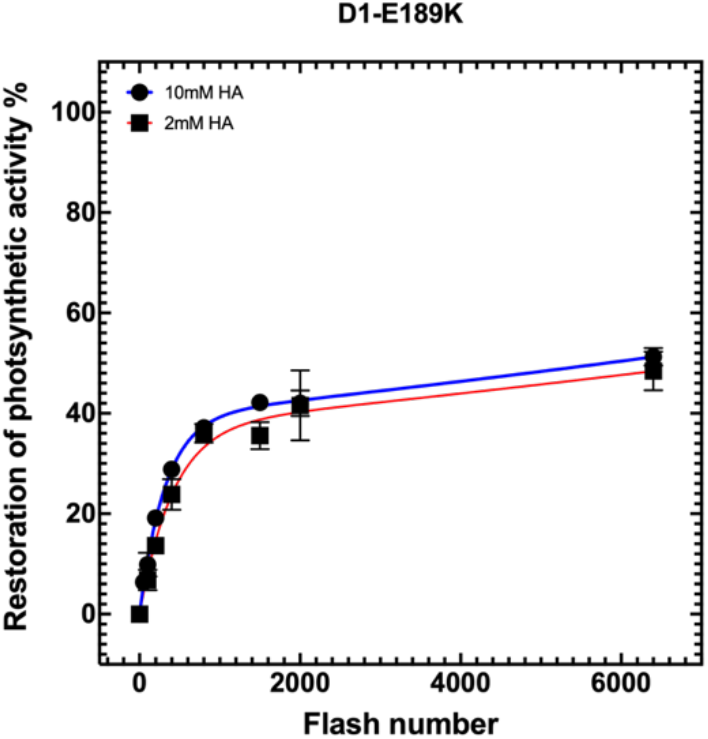
Photoactivation of HA-extracted cells as a function of the single-turnover flash number. Series of xenon lamp flashes were given at uniform frequency of 2 Hz (500ms flash interval) to D1-E189K treated with 2mM and 10mM hydroxylamine. The percentage of O_2_ evolution recovery was calculated as a fraction from the O_2_ evolution rate prior HA extraction 544 ± 16 μmol of O_2_ (mg Chl)^−1^ h^−1^, Error bars represent SD with n ≥ 3.

**Table S3.** Values of the variable fluorescence of the from WT control, D1-D170E, D1-E189K, D1-E189R and D1-E189Q cells prior the HA extraction.

| Strain | Fv ($F_m - F_0/F_0$) |
| --- | --- |
| WT control | 0.78 |
| D1-D170E | 0.65 |
| D1-E189K | 0.65 |
| D1-E189R | 0.44 |
| D1-E189Q | 0.52 |

**Table S4.** Characteristics of chlorophyll fluorescence relaxation kinetics in partially photoactivated, D1-D170E, D1-E189Q and D1-E189R cells.

| Partially photoactivated cells in the presence of DCMU |  |  |  |  |  |
| --- | --- | --- | --- | --- | --- |
| Strain/State | | Fast phase: $\tau$/ms<br>(amplitude (%)) | Middle phase: $\tau$/ms<br>(amplitude (%)) | Slow phase: $\tau$/s<br>(amplitude (%)) | R |
| D1-D170E | Extracted | 6.5 (39.5) | 446 (27.9) | > 100 (32.5) | 0.99 |
|  | Partial Photoactivation | 6 (36.9) | 260 (27.9) | 17.4 (35.3) | 0.99 |
|  | Dark incubated | 9.4 (40.7) | 570 (18.7) | 40.9 (38.9) | 0.99 |
| D1-E189R | Extracted | 2.9 (33.7) | 44 (35.1) | > 100 (32.6) | 0.99 |
|  | Partial Photoactivation | 4.9 (43.3) | 144 (24.2) | 91 (32.4) | 0.99 |
|  | Dark incubated | 1.76 (40.4) | 36 (27.7) | > 100 (31.9) | 0.99 |
| D1-E189Q | Extracted | 0.27 (72.8) | 15.8 (6.5) | > 100 (20.7) | 0.98 |
|  | Partial Photoactivation | 6.2 (26.4) | 551 (16.8) | > 100 (56.9) | 0.99 |
|  | Dark incubated | 7.4 (27.2) | 760 (14.3) | > 100 (58.6) | 0.99 |

## References

1. G. T. Babcock et al., Water oxidation in photosystem II: From radical chemistry to multielectron chemistry. Biochemistry 28, 9557–9565 (1989).

2. T. Cardona, A. Sedoud, N. Cox, A. W. Rutherford, Charge separation in Photosystem II: A comparative and evolutionary overview. Biochimica et Biophysica Acta (BBA) - Bioenergetics 1817, 26–43 (2012).

3. B. Kok, B. Forbush, M. McGloin, Cooperation of charges in photosynthetic O_2_ evolution-I. A linear four step mechanism. Photochem Photobiol 11, 457–475 (1970).

4. H. Bao, R. L. Burnap, Photoactivation: The light-driven assembly of the water oxidation complex of photosystem II. Frontiers in Plant Science 7, 578 (2016).

5. T. A. Ono, H. Mino, Unique binding site for Mn^2+^ ion responsible for reducing an oxidized Y_Z_ tyrosine in manganese-depleted photosystem II membranes. Biochemistry 38, 8778–8785 (1999).

6. G. M. Ananyev, A. Murphy, Y. Abe, G. C. Dismukes, Remarkable affinity and selectivity for Cs+ and uranyl (UO22+) binding to the manganese site of the apo-water oxidation complex of photosystem II. Biochemistry 38, 7200–7209 (1999).

7. R. J. Boerner, A. P. Nguyen, B. A. Barry, R. J. Debus, Evidence from directed mutagenesis that aspartate 170 of the D1 polypeptide influences the assembly and-or stability of the manganese cluster in the photosynthetic water-splitting complex. Biochemistry 31, 6660–6672 (1992).

8. P. J. Nixon, B. A. Diner, Aspartate 170 of the photosystem II reaction center polypeptide D1 is involved in the assembly of the oxygen evolving manganese cluster. Biochemistry 31, 942–948 (1992).

9. P. J. Nixon, J. T. Trost, B. A. Diner, Role of the carboxy terminus of polypeptide-D1 in the assembly of a functional water-oxidizing manganese cluster in photosystem-II of the cyanobacterium *Synechocystis sp* PCC6803 - assembly requires a free carboxyl group at C-terminal position 344. Biochemistry 31, 10859–10871 (1992).

10. H. A. Chu, A. P. Nguyen, R. J. Debus, Site-directed Photosystem-II mutants with perturbed oxygen-evolving properties .2. Increased binding or photooxidation of manganese in the absence of the extrinsic 33-kda polypeptide in-vivo. Biochemistry 33, 6150–6157 (1994).

11. R. O. Cohen, P. J. Nixon, B. A. Diner, Participation of the C-terminal region of the D1-polypeptide in the first steps in the assembly of the Mn4Ca cluster of photosystem II. Journal of Biological Chemistry 282, 7209–7218 (2007).

12. G. M. Ananyev, G. C. Dismukes, High-resolution kinetic studies of the reassembly of the tetra-manganese cluster of photosynthetic water oxidation: proton equilibrium, cations, and electrostatics. Biochemistry 35, 14608–14617 (1996).

13. P. J. Nixon, B. A. Diner, Analysis of water-oxidation mutants constructed in the cyanobacterium *Synechocystis sp*. PCC6803. Biochem Soc Trans 22, 338–343 (1994).

14. Y. Umena, K. Kawakami, J. R. Shen, N. Kamiya, Crystal structure of oxygen-evolving photosystem II at a resolution of 1.9 Ångstrom. Nature 473, 55–U65 (2011).

15. C. J. Gisriel et al., Cryo-EM structure of monomeric photosystem II from *Synechocystis sp* PCC 6803 lacking the water-oxidation complex. Joule 4, 2131–2148 (2020).

16. J. Zabret et al., Structural insights into photosystem II assembly. Nat Plants 7, 524–538 (2021).

17. S. Nakamura, T. Noguchi, Initial Mn2+ binding site in photoassembly of the water-oxidizing Mn4CaO5 cluster in photosystem II as studied by quantum mechanics/molecular mechanics calculations. Chemical Physics Letters 721, 62–67 (2019).

18. C. G. Chen, J. Kazimir, G. M. Cheniae, Calcium modulates the photoassembly of photosystem-II (MN)(4)-cluster by preventing ligation of nonfunctional high-valency states of manganese. Biochemistry 34, 13511–13526 (1995).

19. A. P. Avramov, H. J. Hwang, R. L. Burnap, The role of Ca^2+^ and protein scaffolding in the formation of nature’s water oxidizing complex. Proceedings of the National Academy of Sciences 10.1073/pnas.2011315117, 202011315 (2020).

20. R. J. Debus, K. A. Campbell, D. P. Pham, A.-M. A. Hays, R. D. Britt, Glutamate 189 of the D1 polypeptide modulates the magnetic and redox properties of the manganese cluster and tyrosine Y_Z_ in photosystem II. Biochemistry 39, 6275–6287 (2000).

21. H. A. Chu, A. P. Nguyen, R. J. Debus, Amino-acid-residues that influence the binding of manganese or calcium to photosystem-II.1. The lumenal intehelical domains of the D1 polypepride. Biochemistry 34, 5839–5858 (1995).

22. C. Tommos, G. T. Babcock, Proton and hydrogen currents in photosynthetic water oxidation. Biochimica et Biophysica Acta (BBA) - Bioenergetics 1458, 199–219 (2000).

23. C. Tommos, G. T. Babcock, Oxygen production in nature: A light-driven metalloradical enzyme process. Accounts of Chemical Research 31, 18–25 (1998).

24. H. A. Chu, A. P. Nguyen, R. J. Debus, Amino-acid-residues that influence the binding of manganese or calcium to photosystem-II.2. The carboxy-terminal domain of the D1 polypeptide. Biochemistry 34, 5859–5882 (1995).

25. M. Ibrahim et al., Untangling the sequence of events during the S2 --> S3 transition in photosystem II and implications for the water oxidation mechanism. Proc Natl Acad Sci USA 117, 12624–12635 (2020).

26. M. Suga et al., Light-induced structural changes and the site of O=O bond formation in PSII caught by XFEL. Nature 10.1038/nature21400 (2017).

27. P. E. Siegbahn, Water oxidation mechanism in photosystem II, including oxidations, proton release pathways, O-O bond formation and O_2_ release. Biochimica et biophysica acta 1827, 1003–1019 (2013).

28. N. Cox et al., Electronic structure of the oxygen-evolving complex in photosystem II prior to O-O bond formation. Science 345, 804–808 (2014).

29. Y. Kanesaki et al., Identification of substrain-specific mutations by massively parallel whole-genome resequencing of *Synechocystis sp* PCC 6803. DNA research : an international journal for rapid publication of reports on genes and genomes 19, 67–79 (2012).

30. R. J. Debus et al., Does histidine 332 of the D1 polypeptide ligate the manganese cluster in photosystem II? An electron spin echo envelope modulation study. Biochemistry 40, 3690–3699 (2001).

31. H. A. Chu, A. P. Nguyen, R. J. Debus, Site-directed Photosystem-ii mutants with perturbed oxygen-evolving properties .1. Instability or inefficient assembly of the manganese cluster in-vivo. Biochemistry 33, 6137–6149 (1994).

32. J. G. K. Williams, “Construction of specific mutations in photosystem II photosynthetic reaction center by genetic engineering methods in *Synechocystis 6803*” in Methods in Enzymology. (Academic Press, 1988), vol. 167, pp. 766–778.

33. G. M. Cheniae, I. F. Martin, Effect of hydroxylamine on Photosystem-II .1. Factors affecting decay of O2 evolution. Plant Physiol. 47, 568–& (1971).

34. H. J. Hwang, R. L. Burnap, Multiflash experiments reveal a new kinetic phase of photosystem II manganese cluster assembly in *Synechocystis sp*. PCC6803 *in vivo*. Biochemistry 44, 9766–9774 (2005).

35. H. J. Hwang, A. McLain, R. J. Debus, R. L. Bumap, Photoassembly of the manganese cluster in mutants perturbed in the high affinity Mn-binding site of the H2O-oxidation complex of photosystem II. Biochemistry 46, 13648–13657 (2007).

36. R. Radmer, G. M. Cheniae, Photoactivation of manganese catalyst of O_2_ evolution .2. 2-Quantum mechanism. Biochimica et biophysica acta 253, 182–& (1971).

37. A. P. Avramov, H. J. Hwang, R. L. Burnap, The role of Ca^2+^ and protein scaffolding in the formation of nature’s water oxidizing complex. Proceedings of the National Academy of Sciences 117, 28036 (2020).

38. A. F. Miller, G. W. Brudvig, Manganese and calcium requirements for reconstitution of oxygen evolution activity in manganese-depleted photosystem II membranes. Biochemistry 28, 8181–8190 (1989).

39. N. Tamura, G. Cheniae, Photoactivation of the water-oxidizing complex in photosystem II membranes depleted of Mn and extrinsic proteins. I. Biochemical and kinetic characterization. Biochimica et biophysica acta 890, 179–194 (1987).

40. I. Vass, D. Kirilovsky, A.-L. Etienne, UV-B radiation-induced donor- and acceptor-side modifications of photosystem II in the cyanobacterium *Synechocystis sp*. PCC 6803. Biochemistry 38, 12786–12794 (1999).

41. K. Cser, B. A. Diner, P. J. Nixon, I. Vass, The role of D1-Ala344 in charge stabilization and recombination in photosystem II. Photochemical & photobiological sciences : Official journal of the European Photochemistry Association and the European Society for Photobiology 4, 1049–1054 (2005).

42. R. L. Burnap, M. Qian, C. Pierce, The manganese-stabilizing protein of photosystem II modifies the in vivo deactivation and photoactivation kinetics of the H2O oxidation complex in *Synechocystis sp* PCC6803. Biochemistry 35, 874–882 (1996).

43. P. Chernev et al., Light-driven formation of manganese oxide by today’s photosystem II supports evolutionarily ancient manganese-oxidizing photosynthesis. Nat Commun 11, 6110 (2020).

44. B.-D. Hsu, J.-Y. Lee, R.-L. Pan, The high-affinity binding site for manganese on the oxidizing side of Photosystem II. Biochimica et Biophysica Acta (BBA) - Bioenergetics 890, 89–96 (1987).

45. C. W. Hoganson, D. F. Ghanotakis, G. T. Babcock, C. F. Yocum, Manganese ion reduces redox activated tyrosine in manganese-depleted photosystem II preparations. Photosynth Res 22, 285–294 (1989).

46. B. K. Semin, M. Seibert, A carboxylic residue at the high-affinity, Mn-binding site participates in the binding of iron cations that block the site. Biochim Biophys Acta 1757, 189–197 (2006).

47. B. K. Semin, M. Seibert, Flash-induced blocking of the high-affinity manganese-binding site in photosystem II by iron cations: dependence on the dark interval between flashes and binary oscillations of fluorescence yield. J Phys Chem B 110, 25532–25542 (2006).

48. D. J. Vinyard et al., Photosystem II oxygen-evolving complex photoassembly displays an inverse H/D solvent isotope effect under chloride-limiting conditions. Proceedings of the National Academy of Sciences 116, 18917 (2019).

49. C. A. Tracewell, A. Cua, D. H. Stewart, D. F. Bocian, G. W. Brudvig, Characterization of Carotenoid and Chlorophyll Photooxidation in Photosystem II. Biochemistry 40, 193–203 (2001).

50. T. A. Ono, Y. Inoue, Requirement of divalent cations for photoactivation of the latent water oxidation system in intact chloroplasts from flashed leaves. Biochimica et biophysica acta 723, 191–201 (1983).

51. A. M. Tyryshkin et al., Spectroscopic Evidence for Ca2+ Involvement in the Assembly of the Mn4Ca Cluster in the Photosynthetic Water-Oxidizing Complex. Biochemistry 45, 12876–12889 (2006).

52. D. A. Weisz et al., A novel chlorophyll protein complex in the repair cycle of photosystem II. Proceedings of the National Academy of Sciences 116, 21907 (2019).

53. K. Becker, K. U. Cormann, M. M. Nowaczyk, Assembly of the water-oxidizing complex in photosystem II. J Photochem Photobiol B 104, 204–211 (2011).

54. P. J. Nixon, M. Barker, M. Boehm, R. de Vries, J. Komenda, FtsH-mediated repair of the photosystem II complex in response to light stress. J. Exp. Bot. 56, 357–363 (2005).

55. R. Burnap, L. A. Sherman, Deletion mutagenesis in Synechocystis sp. PCC6803 indicates the the Mn-stabilizing protein of photosystem II is not essential for O2 evolution. Biochemistry 30, 440–446 (1991).

56. E. Busch, E. Hohenester, R. Timpl, M. Paulsson, P. Maurer, Calcium Affinity, Cooperativity, and Domain Interactions of Extracellular EF-hands Present in BM-40*. Journal of Biological Chemistry 275, 25508–25515 (2000).

57. M. Amin et al., Proton-coupled electron transfer during the S-state transitions of the oxygen-evolving complex of photosystem II. J Phys Chem B 10.1021/jp510948e (2015).

58. S. Nakamura, T. Noguchi, Quantum mechanics/molecular mechanics simulation of the ligand vibrations of the water-oxidizing Mn4CaO5 cluster in photosystem II. Proceedings of the National Academy of Sciences 113, 12727–12732 (2016).

59. M. Zhang et al., Structural insights into the light-driven auto-assembly process of the water-oxidizing Mn4CaO5-cluster in photosystem II. Elife 6 (2017).

60. H. Bao, R. L. Burnap, Photoactivation: The light-driven assembly of the water oxidation complex of photosystem II. Front Plant Sci 7, 578 (2016).

61. A. M. Tyryshkin et al., Spectroscopic evidence for Ca^2+^ involvement in the assembly of the Mn4Ca cluster in the photosynthetic water-oxidizing complex. Biochemistry 45, 12876–12889 (2006).

62. C. Chen, J. Kazimir, G. M. Cheniae, Calcium modulates the photoassembly of photosystem II (Mn)4 clusters by preventing ligation of nonfunctional high valency states of manganese. Biochemistry 34, 13511–13526 (1995).

63. G. M. Cheniae, I. F. Martin, Photoactivation of the manganese catalyst of O_2_ evolution. I. Biochemical and kinetic aspects. Biochimica et biophysica acta 253, 167–181 (1971).

64. T. Tokano, Y. Kato, S. Sugiyama, T. Uchihashi, T. Noguchi, Structural Dynamics of a Protein Domain Relevant to the Water-Oxidizing Complex in Photosystem II as Visualized by High-Speed Atomic Force Microscopy. The Journal of Physical Chemistry B 10.1021/acs.jpcb.0c03892 (2020).

65. G. M. Ananyev, G. C. Dismukes, Assembly of the tetra-Mn site of photosynthetic water oxidation by photoactivation: Mn stoichiometry and detection of a new intermediate. Biochemistry 35, 4102–4109 (1996).

66. T.-A. Ono, Y. Inoue, Reductant-sensitive intermediates involved in multi-quantum process of photoactivation of latent O_2_-evolving system. Plant and Cell Physiology 28, 1293–1299 (1987).

67. B. A. Diner, P. J. Nixon, The rate of reduction of oxidized redox-active tyrosine, Z+, by exogenous Mn2+ is slowed in a site-directed mutant, at aspartate 170 of polypeptide D1 of photosystem II, inactive for photosynthetic oxygen evolution. Biochimica et Biophysica Acta (BBA) - Bioenergetics 1101, 134–138 (1992).

